# Same-Slide Spatial Multi-Omics Integration Reveals Tumor Virus-Linked Spatial Reorganization of the Tumor Microenvironment

**DOI:** 10.1101/2024.12.20.629650

**Authors:** Yao Yu Yeo, Yuzhou Chang, Huaying Qiu, Stephanie Pei Tung Yiu, Hendrik A Michel, Wenrui Wu, Xiaojie Jin, Shoko Kure, Lindsay Parmelee, Shuli Luo, Precious Cramer, Jia Le Lee, Yang Wang, Jason Yeung, Nourhan El Ahmar, Berkay Simsek, Razan Mohanna, McKayla Van Orden, Wesley Lu, Kenneth J Livak, Shuqiang Li, Jahanbanoo Shahryari, Leandra Kingsley, Reem N Al-Humadi, Sahar Nasr, Dingani Nkosi, Sam Sadigh, Philip Rock, Leonie Frauenfeld, Louisa Kaufmann, Bokai Zhu, Ankit Basak, Nagendra Dhanikonda, Chi Ngai Chan, Jordan Krull, Ye Won Cho, Chia-Yu Chen, Jia Ying Joey Lee, Hongbo Wang, Bo Zhao, Lit-Hsin Loo, David M Kim, Vassiliki Boussiotis, Baochun Zhang, Alex K Shalek, Brooke Howitt, Sabina Signoretti, Christian M Schürch, F Stephan Hodi, W Richard Burack, Scott J Rodig, Qin Ma, Sizun Jiang

**Author notes:** Co-first authors (Yao Yu Yeo, Yuzhou Chang, Huaying Qiu). Co-second authors (Stephanie Pei Tung Yiu, Hendrik A Michel). Senior authors (Qin Ma, Sizun Jiang).

## Abstract

The advent of spatial transcriptomics and spatial proteomics have enabled profound insights into tissue organization to provide systems-level understanding of diseases. Both technologies currently remain largely independent, and emerging same slide spatial multi-omics approaches are generally limited in plex, spatial resolution, and analytical approaches. We introduce IN-situ DEtailed Phenotyping To High-resolution transcriptomics (IN-DEPTH), a streamlined and resource-effective approach compatible with various spatial platforms. This iterative approach first entails single-cell spatial proteomics and rapid analysis to guide subsequent spatial transcriptomics capture on the same slide without loss in RNA signal. To enable multi-modal insights not possible with current approaches, we introduce k-bandlimited Spectral Graph Cross-Correlation (SGCC) for integrative spatial multi-omics analysis. Application of IN-DEPTH and SGCC on lymphoid tissues demonstrated precise single-cell phenotyping and cell-type specific transcriptome capture, and accurately resolved the local and global transcriptome changes associated with the cellular organization of germinal centers. We then implemented IN-DEPTH and SGCC to dissect the tumor microenvironment (TME) of Epstein-Barr Virus (EBV)-positive and EBV-negative diffuse large B-cell lymphoma (DLBCL). Our results identified a key tumor-macrophage-CD4 T-cell immunomodulatory axis differently regulated between EBV-positive and EBV-negative DLBCL, and its central role in coordinating immune dysfunction and suppression. IN-DEPTH enables scalable, resource-efficient, and comprehensive spatial multi-omics dissection of tissues to advance clinically relevant discoveries.

## Introduction

Spatial transcriptomics and spatial proteomics are recent technological breakthroughs that have enabled investigations of complex biological systems at unprecedented detail within native tissue contexts (1–4). Effective combination of both approaches on the same tissue section is currently the rate-limiting step for novel biological insights, particularly given the complementary strengths of assessing both RNA and proteins. While spatial transcriptomics offers higher feature coverage and pathway-level insights, the technology faces inherent biological limitations in predicting functional outcomes due to post-transcriptional regulation and variable RNA-to-protein correlations (5–7), whereas spatial proteomics directly captures functional molecular phenotypes and functional states with high signal-to-noise ratios and data acquisition speeds, albeit with lower multiplexing capacity. Spatial multi-omics methods that can simultaneously profile both transcripts and proteins from the same tissue section would enable insights into regulatory mechanisms while preserving spatial context to bridge the gap between gene expression and functional protein dynamics in complex biological systems and archival clinical specimens.

Several innovative approaches have successfully demonstrated the potential of integrating spatial protein and RNA imaging on the same tissue sample (8–14). While these pioneering methods have provided valuable insights, current technical constraints, such as multiplexing capacity (8, 10, 11, 14, 15) and spatial resolution in grid/spot-based approaches (8, 9, 12, 13), suggest opportunities for further advancements. Spatial transcriptomics approaches also often incorporate protease treatment of tissue sections for efficient RNA detection, which will compromise protein epitope integrity and impact downstream protein analysis (10, 15, 16). An additional key limitation for broad clinical application and adoption is the compatibility with formalin-fixed paraffin-embedded (FFPE) tissues, the standard preservation method in clinical pathology (17). There is also significant potential to expand computational approaches to fully empower multi-modal analysis for meaningful biological insights (18).

We herein present IN-DEPTH (IN-situ DEtailed Phenotyping To High-resolution transcriptomics), a cost-efficient and reproducible spatial multi-omics approach that utilizes single-cell spatial proteomics to guide subsequent genome-wide spatial transcriptomics capture on the same slide without compromise to protein or RNA signals. IN-DEPTH advances our conceptual approach of spatial multi-omics data generation by linking rapid cell type functional identification and tissue architecture analysis with deep interrogation of transcriptomic pathways in a biologically relevant manner. To quantify tissue spatially-linked transcriptomic pathways revealed by IN-DEPTH, we developed k-bandlimited Spectral Graph Cross-Correlation (SGCC) to determine spatial co-varying relationships between cell pairs using an unbiased graph signal representation method (19). Here, the spatial arrangement and pattern of each cell phenotype is a graph signal where cells serve as nodes, spatial patterns are node attributes, and spatial distances are edges. This allows an unbiased representation of spatial patterns of each cell population on tissues through spectral graph signals to resolve underlying spatial relationships between cell types and gene programs.

We demonstrate the broad applicability of IN-DEPTH across various commercially available spatial platforms, and highlight the combination of IN-DEPTH and SGCC to accurately identify human tonsil multi-modal features at global and local scales. We further demonstrate the synergistic potential of IN-DEPTH and SGCC to unravel novel biological insights on the impact of the prototypic tumor virus, Epstein-Barr Virus (EBV), on the diffuse large B-cell lymphoma (DLBCL) tumor microenvironment (TME) and immune dysregulation. Through our same-slide iterative and integrative spatial multi-omics analysis, we uncover viral-linked spatial reorganization of the DLBCL TME by exploiting a key tumor-macrophage-CD4 T cell immunomodulatory axis to promote CD4 T cell dysfunction, potentially underscoring the need for informed targeted therapeutic strategies in virus-associated malignancies.

## Results

### IN-DEPTH combines antibody staining and RNA probe hybridization on the same slide while retaining protein and RNA quality

IN-DEPTH utilizes high-dimensional spatial proteomics for initial precise cellular phenotyping and functional assessment to guide subsequent targeted spatial transcriptomics capture in specific cell types and regions of interest on the same slide (**Fig. 1A**). This streamlined approach ensures the biological relevance of spatial transcriptomics by tying it to spatial proteomics-guided identification of tissue regions of interest (ROI), thus reducing the resource-intense cost and time barriers associated with spatial transcriptomics of whole slides, while retaining high sensitivity (**Supp Fig. 1A**). Given the impact of the protease digestion step during spatial transcriptomics on subsequent antibody imaging (10, 15, 16), we postulated that performing spatial proteomics first before transcriptomics will circumvent this challenge. As various spatial proteomics platforms also differ in recommended tissue retrieval conditions, we first implemented a standardized heat-induced epitope retrieval step at 97°C for 20 min using a pH 9.0 retrieval buffer followed by a 1-hour photobleaching step, optimized across our prior experiments (10, 11, 20, 21).

**Figure 1:**
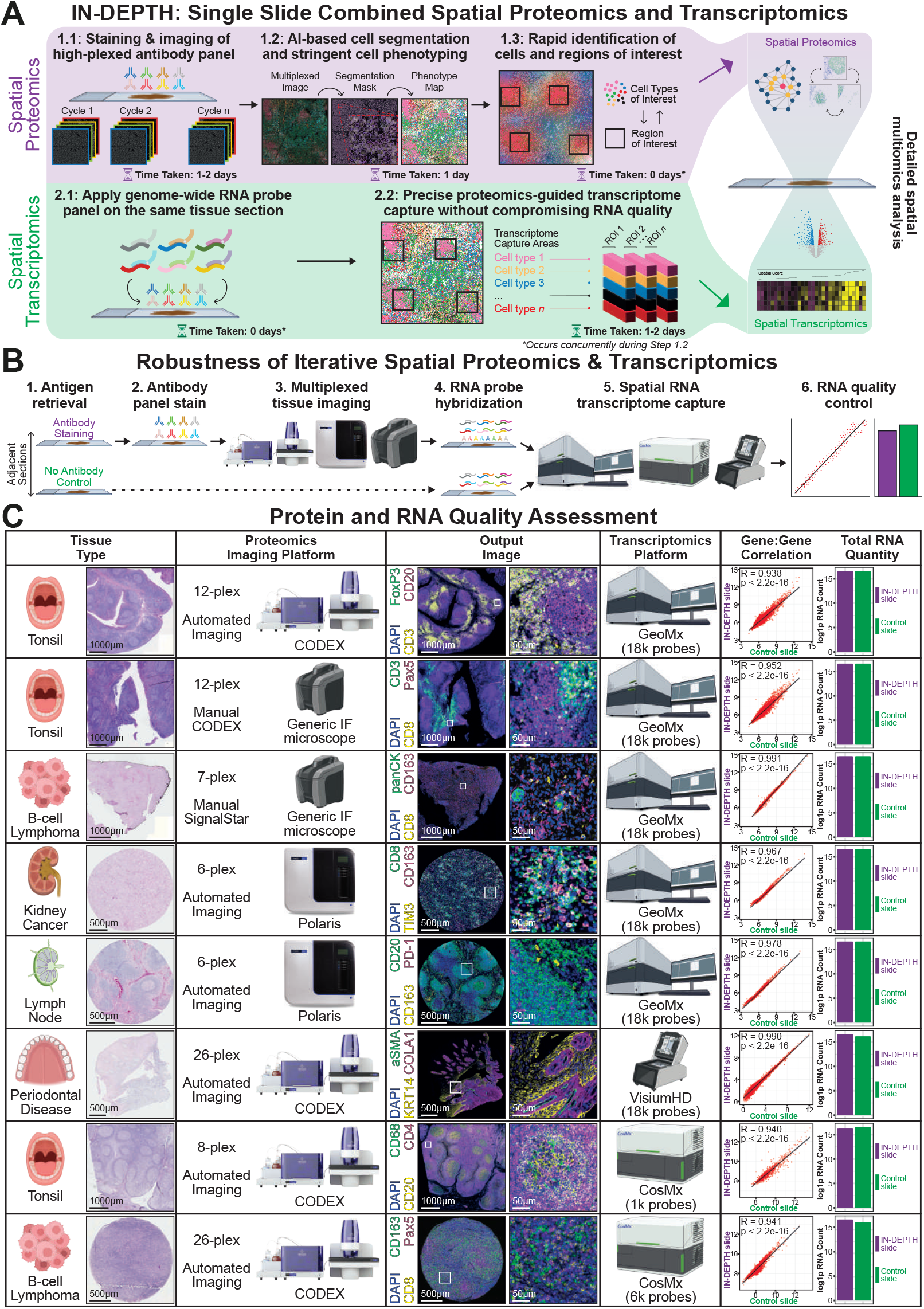
IN-DEPTH combines spatial proteomics and transcriptomics on the same slide without loss of protein or RNA quality. **(A)** Schematic overview of IN-DEPTH, where spatial proteomics was used to guide cell-type specific genome-wide transcriptomic capture on the same slide. **(B)** Experimental outline to assess the effects of spatial proteomics workflow on RNA capture, with an adjacent tissue section without spatial proteomics as a control. **(C)** Assessment of tissue imaging and RNA capture quality after IN-DEPTH. Each row represents a different combination of spatial platforms evaluated for IN-DEPTH and the corresponding tissue type used, and each column represents key experimental variables or data output presented in systematic order from left to right. The breakdown for individual profiled ROIs and negative control probes are in **Supp Figs. 1C & D**. All tissues were subjected to H&E staining at the end of each assay (see **Materials and Methods**).

To systematically evaluate the feasibility of integrating spatial proteomics with transcriptomics with a generalizable framework, we focused on four multiplexed immunofluorescence-based spatial proteomics platforms (CODEX (22), SignalStar (23), Polaris (24), Orion (25)) due to their established track record in clinical applications, general preservation of tissue integrity, rapid whole slide imaging capabilities, and complementary technical approaches to protein labeling. These platforms represent diverse methodologies including cyclic immunofluorescence, signal amplification, and spectral deconvolution, providing a diverse initial setting for method development. We also selected representative spatial transcriptomics platforms (GeoMx (8), VisiumHD (26), CosMx (27)) with broad availability both within and beyond our laboratories, using stringently adjusted protocols to ensure experimental compatibility across both platforms (see **Materials and Methods**).

To determine if prior spatial proteomics on tissue samples affects downstream RNA signal recovery, we first compared the spatial transcriptome signal of adjacent tissue slides, wherein one slide was subjected to IN-DEPTH (spatial proteomics followed by spatial transcriptomics) while the other slide was subjected to only the corresponding spatial transcriptomics platform as a control (**Fig. 1B**). Both slides subsequently underwent hematoxylin and eosin (H&E) staining to assess the retention of tissue morphology. In our initial proof-of-concept, we applied CODEX-GeoMx IN-DEPTH on FFPE tonsil tissues and observed a robust gene-to-gene correlation (R = 0.938) between the IN-DEPTH and the control slide with minimal differences in total captured RNA and robust antibody staining (**Fig. 1C, row 1**). We next demonstrated the easy adaptability of the CODEX approach using any microscope by performing CODEX with manual stripping and hybridization of detection oligos (22, 28) with whole slide imaging using the slide-scanner functionality of the GeoMx instrument followed by RNA recovery, obtaining consistent RNA signals (R = 0.952) (**Fig. 1C, row 2**).

We next expanded upon these initial IN-DEPTH results across various combinations of spatial proteomics and spatial transcriptomics platforms using a variety of FFPE tissue samples. We observed a generally consistent positive gene-to-gene correlation (R > 0.94) and total transcript recovery between the IN-DEPTH and control slides (**Fig. 1C, rows 3-8**), with the exception of the Orion-GeoMx combination with a lower gene-to-gene correlation (R = 0.692) (**Supp Fig. 1B**). The total number of non-binding control RNA probes detected was also consistently low across all conditions (**Supp Fig. 1C**), with the transcriptome gene-to-gene correlation remaining strongly positive across each individual spatially profiled ROI (**Supp Fig. 1D**).

These data collectively demonstrate the robustness of spatial protein and RNA signals with IN-DEPTH, while allowing user flexibility for cross-platform and region-specific RNA capture. Among the validated platform combinations, we selected CODEX-GeoMx for further development based on several key advantages: (1) our strong expertise with the CODEX and GeoMx platforms and experimental protocols compatible with FFPE tissues (10, 11, 20, 21, 29–31), (2) its rapid whole-slide imaging capability enabling comprehensive tissue assessment, (3) access to extensively validated antibody reagents in-house (10, 21, 32, 33) and commercially for tissue profiling, (4) the proven stability and reproducibility in cyclical imaging with CODEX oligo-tagged antibodies (22, 28, 34), and (5) the GeoMx’s ability to automatically capture whole transcriptome data with precise regional selectivity, rapid speed, and cost effectiveness compared to the other transcriptomics platforms we tested (**Supp Fig. 1E**). Based on these advantages, we focused our subsequent IN-DEPTH development and validation on the CODEX-GeoMx platform combination.

### IN-DEPTH enables reproducible and robust spatial multi-omics profiling and reveals functional cell states within the native tissue architecture

We next performed IN-DEPTH (CODEX-GeoMx) on two adjacent FFPE sections from the same tonsil tissue, with each section undergoing RNA capture on two independent GeoMx instruments to assess for technical reproducibility. We applied a 12-plex antibody panel consisting of cell phenotyping markers on both slides together (**Supp Fig. 2A**), and imaged them in parallel on the Phenocycler Fusion system capable of imaging two slides at a time. We performed cell segmentation and phenotyping for 11 cell populations using the background subtracted images from the Phenocycler Fusion (**Fig. 2A, left** and **Fig. 2B, left**).

**Figure 2:**
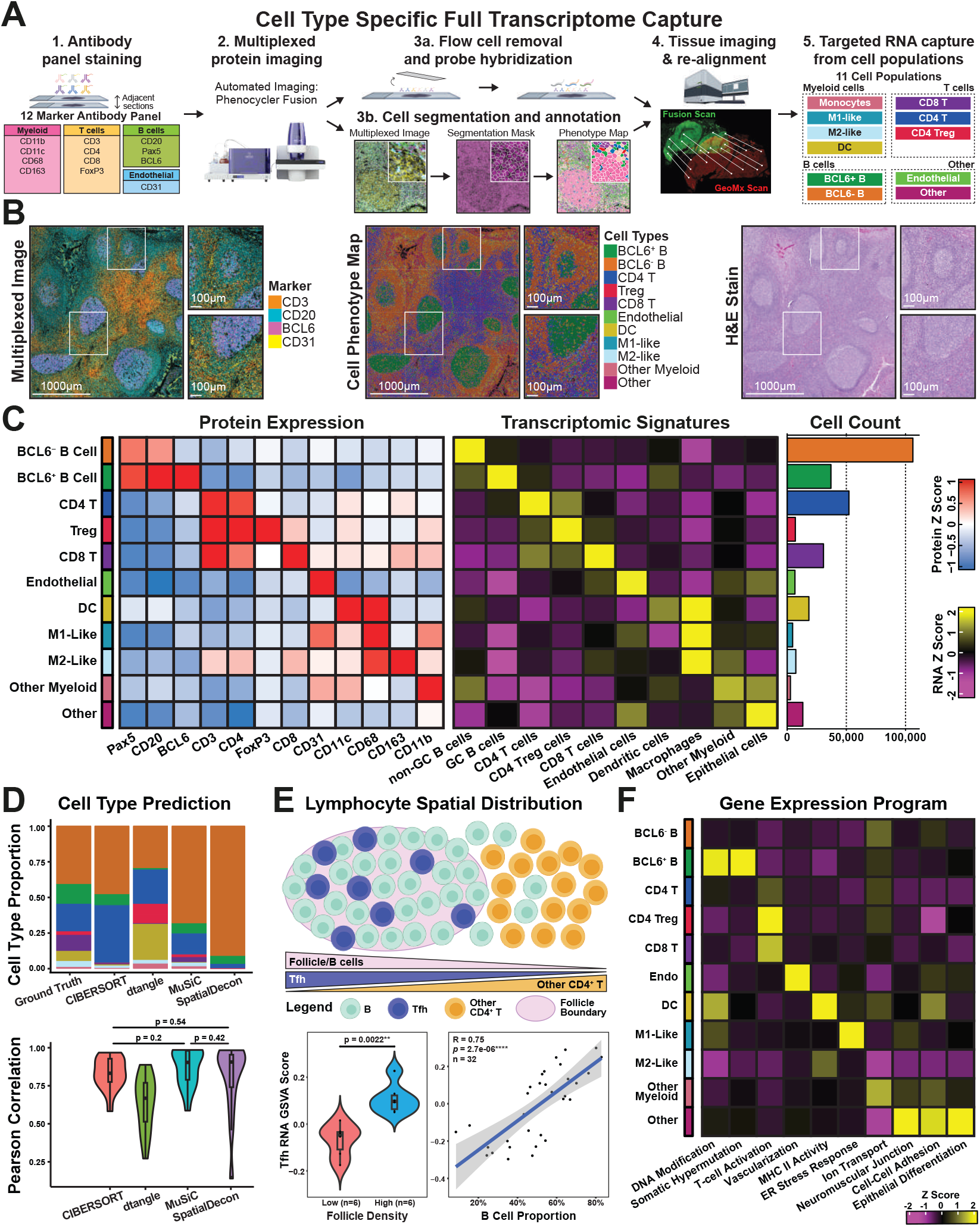
IN-DEPTH enables reproducible and systematic characterization of tonsillar tissue architecture through integrated spatial proteomics and transcriptomics. **(A)** Schematic workflow of IN-DEPTH, illustrating the 12-marker antibody imaging, cell segmentation and phenotyping, cross platform tissue image registration, and targeted RNA capture from identified cell populations on the same slide. **(B)** Visualization of key cellular features in tonsillar tissues using CODEX multiplexed imaging (left) showing T cells (CD3), B cells (CD20 and BCL6), and endothelial cells (CD31), with the corresponding cell phenotype map (middle) and H&E image (right) as part of the IN-DEPTH workflow. **(C)** Cell type-specific protein expression levels (left), gene signatures (middle), and cell counts (right) for the annotated cell types. Data shown is generated from two technical replicates. **(D)** Systematic evaluation of four computational deconvolution algorithms using IN-DEPTH data as the ground truth reference. **(E)** Spatial multi-modal analysistion of Tfh cells showing their distribution relative to B cell follicles (top schematic) and quantitative validation through differential Tfh gene signature enrichment between follicle-high and follicle-low regions (bottom left, 6 ROIs chosen each), and correlation with B cell density (bottom right). A two-sided Wilcoxon rank sum test was performed, with the null hypothesis that there is no difference in the Tfh signature between follicle-low and follicle-high regions (bottom left), and a Spearman’s correlation was used for the correlation test (bottom right). **(F)** Top cell type-specific gene expression programs identified, and their relative enrichment across the 12 annotated cell populations.

To capture cell type-specific transcriptomes, we imported these cell-type specific masks onto the GeoMx for custom spatial transcriptome capture using the human whole transcriptome atlas (hWTA) library consisting of >18,000 targets in the human genome. We selected 16 paired and continuous 660×760 µm rectangular ROIs on each adjacent slide that include B follicles and T cell zones (**Supp Fig. 2B**). We first confirmed the specificity of our antibody panel and accuracy of spatial proteomics cell type annotation for both tissues (**Fig. 2B, middle** and **Supp Fig. 2C, middle**), with final confirmatory assessment with board-certified pathologists by assessing the post-IN-DEPTH H&E staining of the same tissue section (**Fig. 2B, right** and **Supp Fig. 2B, right**).

We further assessed the specificity and accuracy of our cell phenotyping via the expected enriched expression of each antibody marker in each of the 11 annotated cell populations (**Fig. 2C, left** and **Supp Fig. 2D, left**). We then orthogonally verified the spatial transcriptomics capture specificity by quantifying the enrichment of cell-type specific transcriptomic signatures for each cell population against a single-cell tonsil atlas (35) (**Fig. 2C, middle** and **Supp Fig. 2D, middle**). We additionally confirmed the expected cell counts (**Fig. 2C, right** and **Supp Fig. 2D, right**), high consistency between the protein and transcriptome signatures (**Supp Fig. 2D**), gene-to-gene correlation (**Supp Fig. 2E**), total RNA capture (**Supp Fig. 2F**), and low signals from non-targeting negative control probes (**Supp Fig. 2G**) between the adjacent slides. These results highlight the robust technical reproducibility of IN-DEPTH across different instruments.

We recognize that spatial proteomics-guided transcriptomes with IN-DEPTH is well suited to address the challenge of accurate real world ground-truth reference data currently missing for deconvolution approaches (36–39). We demonstrate this application by systematically bench-marking the performances of popular deconvolution algorithms CIBERSORT (40), dtangle (41), MuSic (42), and SpatialDecon (43) on our reference gene signatures (**Supp Table 1** and see **Material and Methods**). We observed that for the top three cell type components — BCL6-positive B cells, BCL6-negative B cells, and CD4 T cells — the results from CIBERSORT, dtangle, and MuSiC were relatively consistent (**Fig. 2D**). Ranking the tonsil ROIs by cell type proportion complexity, as estimated by the Gini-Simpson index (**Supp Fig. 2H**) applied to ground truth cell type proportions, revealed that all four methods achieved high correlation (>0.9) with the ground truth in ROIs of low complexity (e.g. ROIs 1, 2, 3, 4, 5, 9). These results not only validate IN-DEPTH’s ability to generate reliable ground-truth spatial references, but also provide valuable insights for selecting and optimizing computational approaches for specific tissue contexts and research questions.

To demonstrate the utility of paired spatial proteomics and transcriptomics data from IN-DEPTH, we next examined the established functional and spatial dynamics of lymphocytes in the tonsillar tissue architecture. We focused on CD4 T follicular helper (Tfh) cells, which are known to migrate into B follicles during their activation and maturation process (35) (**Fig. 2E, top**). While Tfhs can be easily identified from our CD4 T cell population as spatially residing within B follicles, we did not include Tfh-specific markers such as PD-1 or CXCR5 in our study, making them difficult to annotate using canonical spatial proteomics analysis. We first hypothesized an enrichment in Tfh gene signatures (**Supp Table 1**) for CD4 T cells located in the follicles compared to those outside. Our results confirmed a significant increase in Tfh gene set variation analysis (GSVA) signatures in the ROIs stratified by high or low B follicle densities (**Fig. 2E, bottom left**). We further identified a positive correlation between the Tfh GSVA scores with the proportion of B cells across all ROIs from both tissues (R = 0.75) (**Fig. 2E, bottom right**), consistent with the known Tfh cell trafficking and maturation in the tonsil (35).

To systematically characterize tissue-wide, cell type-specific transcriptional programs in the tonsil, we performed consensus non-negative matrix factorization (44) to infer the predominant gene expression programs (GEPs) within the tonsil for each cell type and identified 10 distinct GEPs. These GEPs were annotated based on Gene Ontology Biological Process (GOBP) signatures (**Supp Table 2**) and exhibited cell type-specific distributions aligning with known cellular functions (35). These specifically include “DNA Modification” and “Somatic Hypermutation” in BCL6-positive B cells, “T cell Activation” across T cells, “Vascularization” in endothelial cells, “MHC Class II Activity” in dendritic cells and M2-like macrophages, “ER Stress Response” in M1-like macrophages, and “Epithelial Differentiation” in Other (non-immune) cells that predominantly reside in tonsillar crypts (**Fig. 2F**).

The reproducible spatial and molecular profiling demonstrated here, from precise cell type identification to capture of transitional cell states and tissue wide transcriptional programs, establishes IN-DEPTH as a robust platform for deep multi-omics investigation of tissue biology. Beyond elucidating detailed cellular and molecular profiles in their native context, IN-DEPTH also enables essential reference data to advance computational approaches such as cell deconvolution.

### Coordinated spatial transitions in cellular states and tissue organization

To investigate how spatial organization relates to cellular function and maximize the utility of IN-DEPTH multiomics data, we developed Spectral graph cross-correlation (SGCC), a mathematical formulation built upon graph signal processing approaches to analyze pairwise coordinated spatial patterns. SGCC leverages the unbiased representation and interpretability of Graph Fourier transform (GFT) to explore the distributional relationships between pairs of cell phenotypes. In our previous study (19), any spatial-omics feature (e.g. cell phenotype labels) can be treated as a graph signal, where the underlying graph can be a lattice graph (a pixel graph with nodes representing pixels and edges defined by pixel-to-pixel distance) or an irregular graph (a cell graph with nodes representing cells and edges defined by cell-to-cell distance). Subsequently, GFT is applied to project vertex-domain graph signals onto the frequency domain via Fourier modes (FM) (see **Materials and Methods**), yielding a set of interpretable Fourier coefficients (FC). As the first k low-frequency FMs capture the spatially organized components of the graph signal (45, 46), it lays the foundation of correlating pairwise cell phenotype in frequency domain by computing the similarity of these k-bandlimited Fourier coefficients.

SGCC quantitatively measures the spatial distributional relationships and underlying patterns between two cell phenotypes via the following three steps. First, by binning cell phenotypes from the cell graph into a pixel graph, all ROIs’ FCs are placed within the same linear space, ensuring subsequent cross-correlation calculations. Second, the binned cell phenotype data are transformed into the frequency domain via Graph Fourier Transform. A low-frequency bandwidth is then delineated, enabling the extraction and selection of the top k band-limited Fourier coefficients that characterize the broad-scale spatial organization. Third, pairwise correlations between cell phenotypes are computed, resulting in c(m,2) pairwise comparisons, where m represents the number of cell phenotypes. These SGCC scores reflect the spatial distribution patterns between two cell types (**Fig. 3A**).

**Figure 3.**
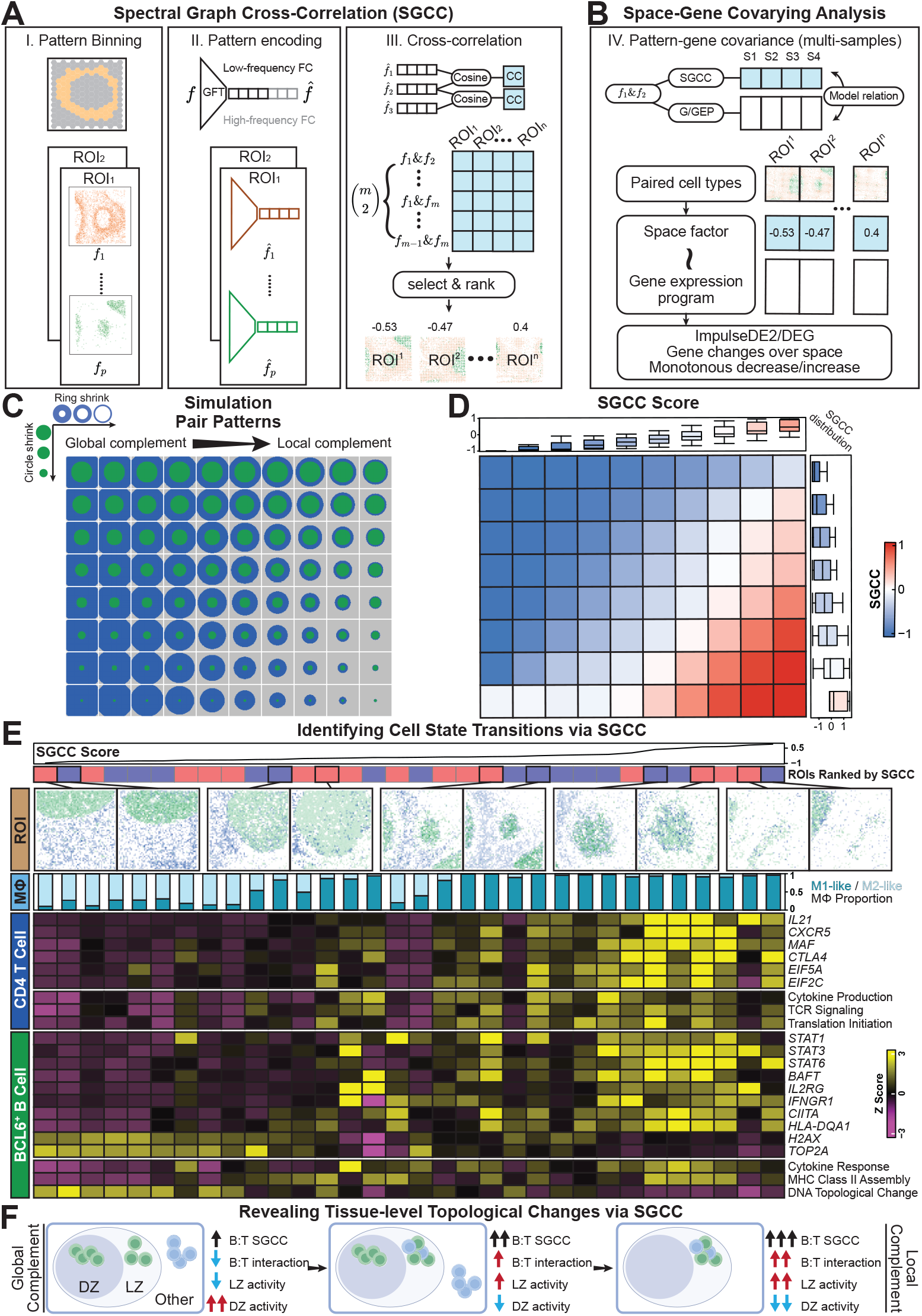
SGCC reveals coordinated spatial transitions in cellular states and tissue architecture. **(A)** Schematic overview of the SGCC methodology showing: I) Pattern binning of single-cells in spatial proteomics data, followed by II) Pattern encoding through GFT to generate low-frequency FCs, and III) Cross-correlation analysis to identified coordinated spatial patterns for downstream integration with transcriptomics. **(B)** Integration framework for identifying genes covarying with spatial pattern across the tissue, linking spatial factors to gene expressionfor functional analysis. **(C)** Systematic validation of SGCC using 80 simulated spatial patterns to demonstrate the ability to detect transitions from global to local complement states. **(D)** Quantification of pattern relationships through SGCC scores. **(E)** Analysis of CD4 T cell and BCL6-positive B cells via IN-DEPTH proteomics and transcriptomics analysis, showing SGCC scores and their associated spatial distribution of cells in bins (top), changes in macrophage polarization states (M1/M2-like proportion), and coordinated gene expression programs reflecting intrinsic cell programs and T-B cell crosstalk (bottom). The full gene pathway names can be found in **Supp Table 2. (F)** A schematic illustrating tissue-level organization derived from SGCC analysis depicting the transitions in T-B cell interactions across the dark zone (DZ) and light zone (LZ).

When multiple samples are available, SGCC can be treated as a continuous or ordinal variable serving as a spatial factor. A negative SGCC value indicates reduced spatial co-occurrence, while a positive value indicates increased spatial co-occurrence between cell phenotypes. Consequently, SGCC can be used to predict genes covarying with spatial factors. For example, one can apply the ImpulseDE2 model (47) to treat SGCC as a continuous spatial variable, or employ edgeR (48) to treat it as an ordinal spatial variable, thereby enabling the identification of spatially dynamic genes (**Fig. 3B**).

We first simulated 80 datasets, each representing a 60×60 pixel graph, to create ring-like distributions of two cell phenotypes. These distributions varied in terms of area and complementarity, thus demonstrating both global and local patterns (**Fig. 3C**). Next, we conducted a k-bandlimited Fourier mode selection experiment to identify the optimal number of neighbors for ensuring robust graph smoothness, thhereby defining the robust low-frequency Fourier modes. As shown in **Supp Figs. 3A & 3B**, when the graph size is 60 nodes and the number of neighbors is set to 400, the graph’s smoothness remains stable at the eigenvalue “knee” point following Laplacian decomposition. We then computed their SGCC scores, which increased under locally complementary patterns but decreased under globally complementary patterns, indicating that SGCC effectively distinguishes changes in spatial patterns (**Fig. 3D**). Another additional set of 80 cell phenotype pixel graphs demonstrated that SGCC can also discriminate differences in area and spatial proximity between two cell phenotype patterns (**Supp Figs. 3C & 3D**).

We next demonstrated the applicability of SGCC to real world IN-DEPTH data, by stratifying nuanced cell state transitions between CD4 T cells and BCL6-positive B cells, key players in modulating germinal center reactions (49). The SGCC score between these cell populations identified consistent orchestrated spatial patterns between tissue replicates (**Supp Figs. 3E & F**). Increasing SGCC between these T and B cells revealed coordinated changes in tissue organization and macrophage cell states (**Fig. 3E, top**), with a more immunosuppressive M2-like polarization toward a more reactive M1-like state as the SGCC score increases, along with a decrease in CD163 expression (**Supp Fig. 3G**). This analysis also uncovered gradual changes in gene expression signatures, reflecting an increase in T cell and B cell cytokine production, B cell MHC-II, T cell TCR activation, and B cell PAX5 expression (**Supp Fig. 3G**) with increased SGCC score and a transition from global to local complementary patterns (**Fig. 3E, bottom**). These transcriptional changes were associated with the functional states of CD4 T cells and follicular B cells, where the low SGCC regions align with self-aggregation of T cells and B cells (**Figs. 3E & F, left**), and the high SGCC regions align with more T-B cell crosstalk akin to light zone interactions (**Figs. 3E & F, right**). These data together demonstrate the unique insights enabled by the combination of IN-DEPTH spatial multi-omics data with SGCC analysis to reveal spatially coordinated transitions in cell states and function, beyond the capacity of either modality alone.

### IN-DEPTH reveals an EBV-linked macrophage immunosuppression and associated CD4 T cell dysfunction in the DLBCL TME

To investigate the complex tumor-immune interactions in the viral-linked TME, we next applied IN-DEPTH to dissect the poorly understood TME of EBV-positive and EBV-negative DLBCL. Using a multi-institutional cohort of FFPE tissues from 17 EBV-positive and 13 EBV-negative patients, we performed IN-DEPTH (CODEX-GeoMx) with a 30-marker antibody panel for cell phenotyping and functional analysis (**Fig. 4A** and **Supp Fig. 4**). We identified 8 distinct cell populations (**Fig. 4A** and **Supp. Fig.5**, from which we captured genome-wide transcriptomes across 38 ROIs (one per patient) with appropriate batch effect correction applied (see **Materials and Methods** and **Supp Fig. 6A**). All images and annotations were validated through same-slide H&E review by board-certified pathologists (**Figs. 4B & C**).

**Figure 4.**
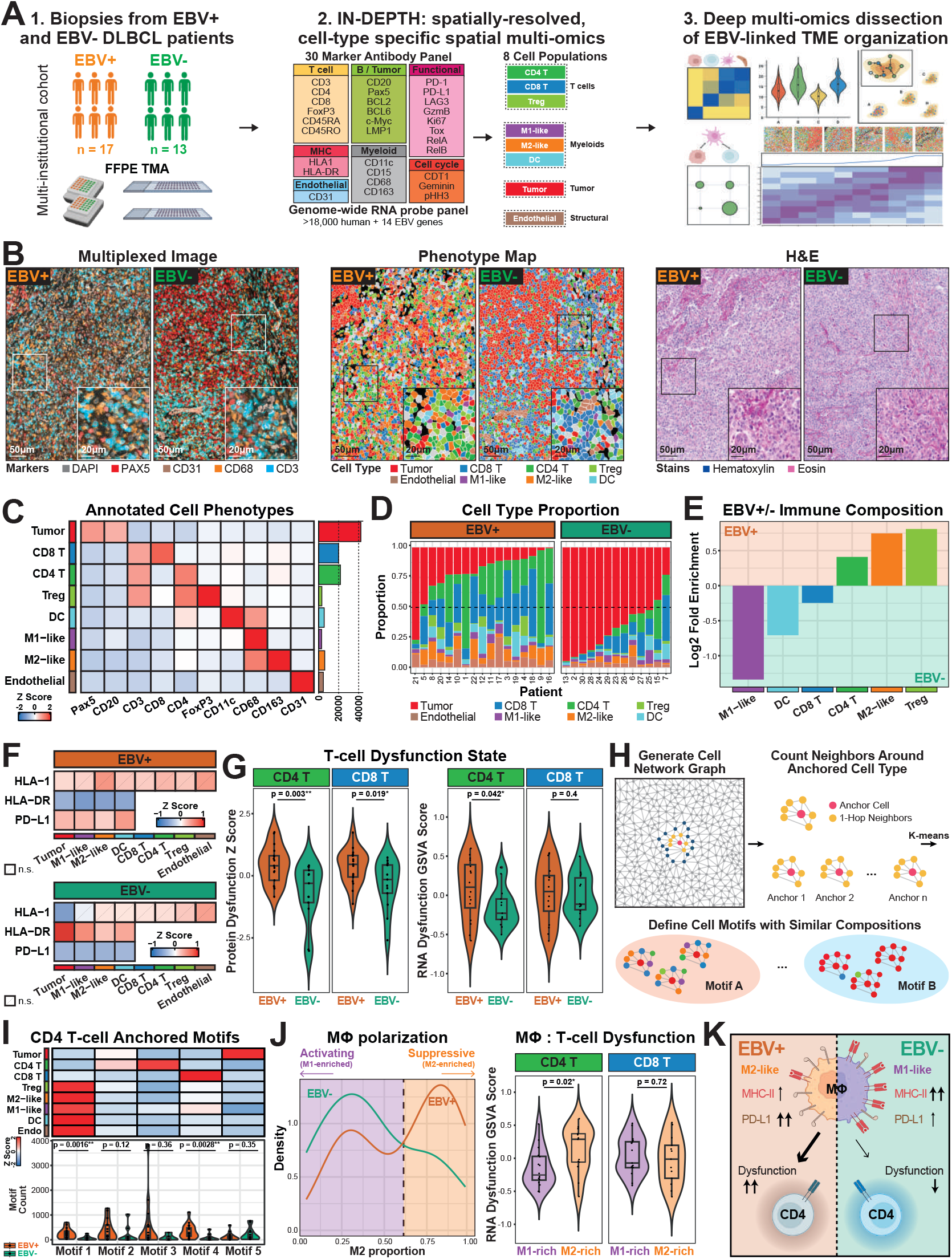
Iterative spatial multi-omics dissection of EBV-positive and EBV-negative DLBCL via IN-DEPTH reveals a macrophage-linked CD4 T cell dysfunction interaction axis. **(A)** IN-DEPTH workflow on EBV-positive (n=17) and EBV-negative (n=13) DLBCL biopsy samples, using a 30-marker antibody panel and a genome-wide RNA probe panel spiked in with custom-designed probes targeting 14 EBV genes. **(B)** Representative CODEX multiplexed images (left) with markers for nuclei (DAPI), B/tumor cells (Pax5), endothelial cells (CD31), macrophages (CD68), and T cells (CD3) shown, as well as the corresponding phenotype maps (middle), and H&E images (right) of EBV-positive and EBV-negative DLBCL tissues. Phenotype maps for each tissue sample core are in **Supp Fig. 5. (C)** Relative protein expression levels (left) and cell counts (right) for the annotated cell types from this DLBCL cohort. **(D)** Relative proportions of annotated cell types across EBV-positive and EBV-negative (left) tissues. **(E)** Log2 fold enrichment plot of immune cell proportions between EBV-positive and EBV-negative DLBCL tissues in this patient cohort. **(F)** Relative protein expression of MHC Class I (HLA1), MHC Class II (HLA-DR), and PD-L1, on the corresponding cell types that express these molecules across EBV-positive (top) and EBV-negative (bottom) DLBCL tissues in this patient cohort. **(G)** Left: Comparison of CD4 and CD8 T cell dysfunction scores calculated based on protein markers between EBV-positive and EBV-negative DLBCL tissues. Right: Comparison of CD4 and CD8 T cell dysfunction scores calculated based on GSVA scoring of RNA signatures EBV-positive and EBV-negative DLBCL tissues. A one-sided Wilcoxon rank sum test were performed, with the alternative hypothesis that the T cell dysfunction signature was greater in the EBV-positive tissues. The protein markers and RNA signatures were curated using a panel of T cell exhaustion checkpoint markers and genes (see **Materials and Methods**). **(H)** Schematic representation of identifying different cellular motifs through n-hop neighborhood analysis anchored on a cell type of interest. **(I)** Top: Cell type enrichment from each identified cellular motif, with CD4 T cells set as the anchor cell. Bottom: Comparison of motif abundance between EBV-positive and EBV-negative DLBCL. A two-sided Wilcoxon rank sum test was performed, with the null hypothesis that there is no difference between motif abundance in EBV-positive and EBV-negative tissues. **(J)** Left: Distribution of the density of M2-like macrophages between EBV-positive and EBV-negative DLBCL tissues in this patient cohort, with the dotted line indicating the cutoff for stratifying M1-rich and M2-rich samples. Right: Comparison of RNA GSVA score of CD4 and CD8 T cell dysfunction between M1-rich and M2-rich populations. A one-sided Wilcoxon rank sum test was performed, with the alternative hypothesis that the T cell dysfunction signature was greater in the EBV-positive tissues. **(K)** Cartoon model depicting key differences in macrophage and CD4 T cell dysfunction states between EBV-positive and EBV-negative DLBCL.

Building upon our prior findings of increased T cell dysfunction in EBV-positive classical Hodgkin’s Lymphoma (cHL) TME (21), we hypothesized there to be distinctive immune composition and organization within the EBV-stratified DLBCL TME. Our initial analysis revealed striking differences in TME composition, with EBV-positive DLBCL consisting of higher immune infiltrates compared to the tumor-heavy EBV-negative cases (**Fig. 4D**). Further dissection of the immune population demonstrated an EBV-associated increase in regulatory T cells (Tregs), and a distinctive shift in macrophage polarization marked by elevated immunosuppressive M2-like macrophages and diminished reactive M1-like macrophages in the EBV-positive DLBCL (**Fig. 4E** and **Supp Fig. 6B**).

At the tissue level, the EBV-positive DLBCL TME exhibited reduced MHC Class II expression, elevated PD-L1, and minimal differences in MHC Class I (**Fig. 4F**), suggesting a CD4 T cell-focused mechanism of dysfunction. Using CD4 and CD8 T cell dysfunction signatures on both the protein and transcript levels (50–52), we found increased global T cell dysfunction in EBV-positive DLBCL, with CD4 T cells exhibiting significantly more pronounced effects than CD8 T cells (**Fig. 4G**). The orthogonal confirmation of T cell dysfunction at both protein and transcript levels highlight the value of same-slide multi-omics via IN-DEPTH for biological discovery and validation.

To identify the cellular neighborhoods associated with elevated CD4 T cell dysfunction in EBV-positive DLBCL, we analyzed the immediate network of cells surrounding CD4 T cells using a network graph approach on the most immediately adjacent (1-hop neighbors). K-means clustering classified 5 distinct motifs (**Fig. 4H** and **Supp Figs. 6C & D**), with immune-rich Motif 1 (enriched in macrophages, Tregs, dendritic cells, and endothelial cells) and 4 (enriched in CD8 T cells) significantly more prevalent in EBV-positive cases and no significant EBV-linked differences for the other motifs (**Fig. 4I**). Further comparison of the protein-derived CD4 T cell dysfunction scores between EBV-positive and EBV-negative immune-enriched (Motifs 1) and immune-deficient motifs (Motifs 2 + 3 + 4 + 5) revealed a graded decrease in CD4 T cell dysfunction from EBV-positive immune-enriched to EBV-negative immune-deficient motifs (**Supp Fig. 6E**).

Given the role of macrophages as major MHC Class II antigen-presenting cells and immune modulators in the TME (53), we examined their contribution to CD4 T cell dysfunction between EBV-positive and EBV-negative DL-BCL. We performed negative binomial regression on M1-like and M2-like macrophages (**Supp Fig. 6F** and **Supp Table 3**), and identified that EBV-positive samples had approximately 1.91 times the expected M2-like macrophage count compared to EBV-negative samples (p < 0.05, 95% confidence interval [1.64, 2.25]) for any given motif. In contrast, the expected M1-like macrophage count compared to EBV-negative samples was 0.86 times that of EBV-positive DLBCL (p < 0.05, 95% confidence interval [0.74, 0.99]). Macrophage association with EBV-negative tumors had decreased PD-L1 and increased HLA-DR with higher tumor density, with both trends reversed in LMP1-positive EBV-positive tumor cells (**Supp Fig. 6G**). These findings implicate a key role of immunosuppressive macrophages as key modulators of CD4 T cell dysfunction in EBV-positive DLBCL. We also observed a clear bimodal distribution of macrophage polarization associated with EBV status, with an elevation of suppressive M2-like in EBV-positive and activating M1-like in EBV-negative cases (**Fig. 4J, left**). Notably, M2-enriched regions displayed significantly higher CD4 T cell dysfunction signatures, with no corresponding differences in CD8 T cell dysfunction (**Fig. 4J, right**). These findings support a model in which EBV reshapes the DLBCL microenvironment through a coordinated reduction in MHC Class II, elevation of PD-L1, and conditioning of an M2-polarized macrophage microenvironment around CD4 T cells to promote T cell dysfunction (**Fig. 4K**).

### SGCC analysis reveals a spatially coordinated tumor– macrophage-CD4 T cell axis driving immune dysfunction in EBV-linked DLBCL

To further dissect the molecular mechanisms underpinning our proposed model of EBV-linked CD4 T cell dysfunction (**Fig. 4K**), we extended SGCC to analyze the spatial relationships between tumor cells, macrophages and CD4 T cells and elucidate coordinated molecular mechanisms underlying this biological process.

First examining tumor-macrophage interactions, we confirmed EBV presence through viral transcript detection and LMP1 viral oncoprotein expression in EBV-positive tumors (**Fig. 5A, top** and **Supp Fig. 7A, top**). As EBV is primarily present in tumor cells, we assessed how tumor cells can influence macrophage functional states. SGCC analysis revealed divergent immunomodulatory signatures: EBV-negative tumor cells exhibited M1-polarizing signatures while EBV-positive tumor cells promoted an immunosuppressive M2-like TME (**Fig. 5A, middle** and **Supp Fig. 7A, middle**). This observation was further supported by the macrophage phenotype distribution and transcriptional programs, showing a predominantly M2-like phenotype and gene program in EBV-positive and M1-like in EBV-negative cases, with increased SGCC scores (**Fig. 5A, bottom** and **Supp Fig. 7A, bottom**). We next assessed the influence of macrophages on CD4 T cell functional states (**Fig. 5B, top**). In EBV-negative DLBCL, increasing SGCC was associated with MHC Class II gene program and HLA-DR protein expression along with T cell activation signatures, which were conversely dampened in EBV-positive DLBCL (**Fig. 5B, middle** and **Supp Fig. 7B, top**). This was consistent with the increase in T cell dysfunction states and low T cell activation pathways in EBV-positive DLBCL, with both trends reversed in EBV-negative DLBCL, as SGCC scores increased (**Fig. 5B, bottom** and **Supp Fig. 7B, bottom**). These findings support a key spatially-linked and immunomodulatory role of macrophages in inducing contrasting CD4 T cell functional states specific for the EBV-positive TME.

**Figure 5.**
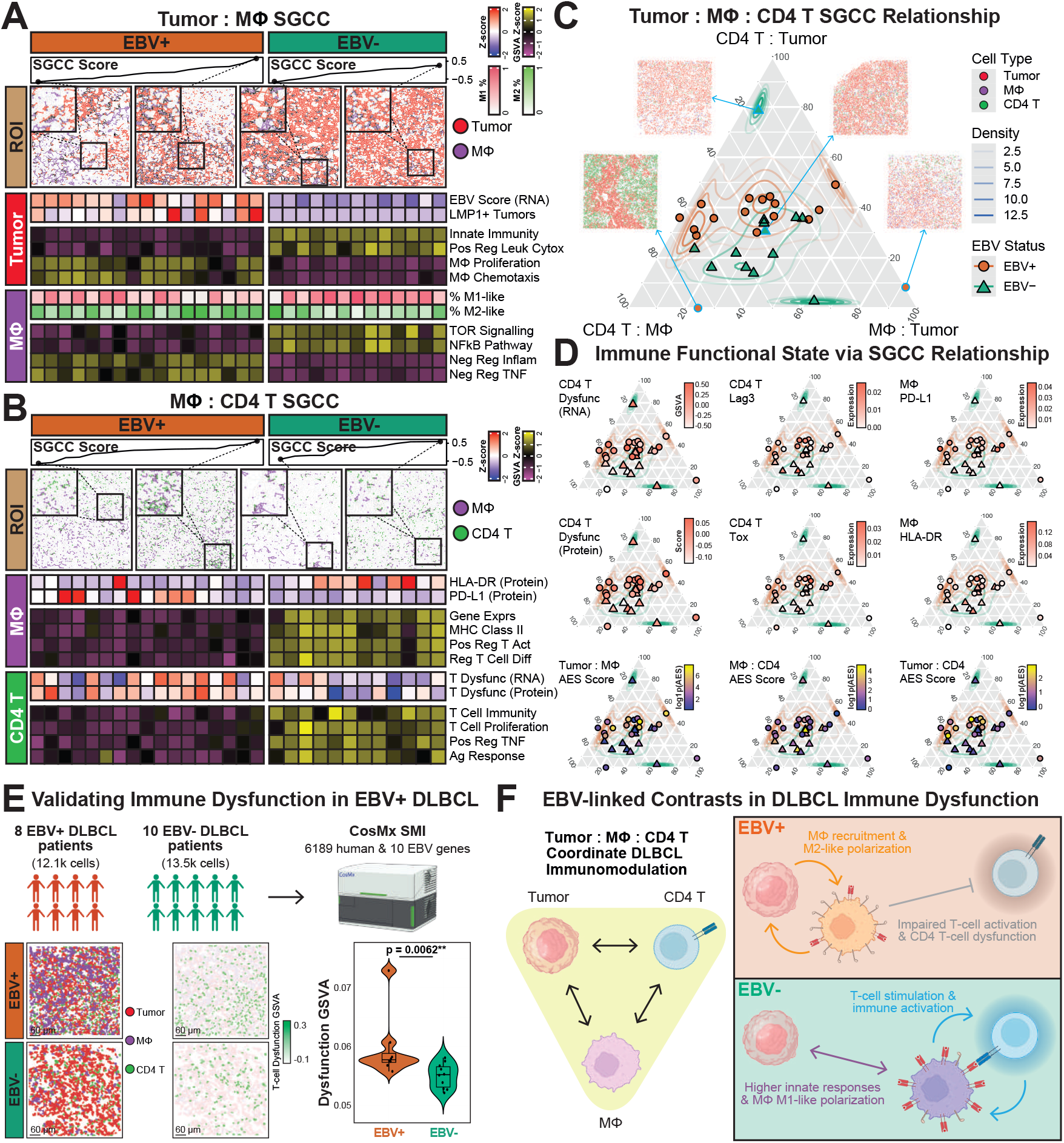
SGCC reveals coordinated spatial multi-modal interactions and EBV-linked cell states in the tumor-macrophage-CD4 T cell axis. **(A)** Analysis of tumor-macrophage spatial relationships. Top: SGCC-ranked spatial distributions and representative images. Middle: EBV transcript levels, LMP1+ tumor cells, and tumor-associated signaling pathways across SGCC scores. Bottom: Changes in macrophage M1/M2 polarization states and associated pathway signatures with increasing SGCC scores. **(B)** Analysis of macrophage-CD4 T cell interactions. Top: SGCC-ranked spatial distributions and representative images. Middle: Changes in PD-L1 and HLA-DR expression of macrophage and antigen presentation pathways across SGCC scores. Bottom: Changes in T cell dysfunction signatures and immune activation pathways across SGCC scores. The full gene pathway names for (A) and (B) are in **Supp Table 2. (C)** Ternary plot depicting a three-way SGCC relationship between CD4 T cells and tumor (top vertex), CD4 T cells and macrophages (bottom left vertex), and macrophages and tumor (bottom right vertex). Points located near the vertices indicate colocalization between two specific cell types while forming a complementary structure with the third cell type (e.g. the ROI from Rochester 4 at the left bottom end of the triangle demonstrates colocalization between CD4 T cells and macrophages while complementing the tumor). In contrast, points near the center of the triangle may signify colocalization among all three cell types. **(D)** Ternary plots across the tumor-macrophage-CD4 T cell axis colored by their expression of key immune dysfunction features (top two rows) or adjacency enrichment statistic (AES) (bottom row). **(E)** Validation in an independent cohort using CosMx. Top: Study design with EBV-positive (n=8) and EBV-negative (n=10) DLBCL biopsy samples using a 6k-plex panel. Bottom left: Representative phenotype map of one EBV-positive and one EBV-negative FOV, showing the spatial organization of annotated tumor (red), macrophage (purple), and T cell populations (green). Bottom middle: Re-visualizing the same phenotype map to emphasize T cell dysfunction GSVA score on T cells. Bottom right: Comparison of T cell dysfunction GSVA scores between EBV-positive and EBV-negative tissues from this cohort. A two-sided Wilcoxon rank sum test was performed, with the null hypothesis that there is no difference in T cell dysfunction score between EBV-positive and EBV-negative tissues. **(F)** Cartoon model depicting contrasting immune state differences in the tumor-macrophage-CD4 T cell interaction axis between EBV-positive (more immunosuppressive) and EBV-negative (less immunosuppresive) DLBCL TMEs.

To better appreciate the complexities of this tripartite spatial interaction, we visualized three-way relationships using ternary analysis of SGCC scores (**Fig. 5C**). While SGCC scores were generally evenly distributed between these 3 cell populations, we observed an enrichment of CD4 T cell-centric SGCC scores in EBV-positive, and macrophage-centric scores in EBV-negative DLBCL TMEs. Markers of T cell dysfunction peaked where all three cell types co-localized (**Fig. 5D, rows 1-2** and **Supp Table 4**), indicative of this tripartite spatial interaction axis in promoting CD4 T cell dysfunction. Adjacency enrichment statistic (AES) analysis (54) further revealed preferential tumor-macrophage interactions in EBV-positive DL-BCL versus macrophage-CD4 T cell interactions in EBV-negative cases (**Fig. 5D, row 3** and **Supp Table 4**), supporting a model in which tumor-macrophage crosstalk and immunosuppression predominates in the EBV-positive DLBCL TME to limit CD4 T cell activation and promote dysfunction. LMP1 appears to play a role here, with an enrichment in expression at the center of the ternary plot (**Supp Fig. 7C**) and positive correlation of LMP1-expressing tumor cells with M2-like polarization and CD4 T cell dysfunction (**Supp Fig. 7D**). Conversely, macrophage-mediated CD4 T cell engagement is more prevalent in EBV-negative DLBCL, facilitating an immune reactive TME with increased CD4 T cell activation and functional immune responses.

We validated our observations using a 6k-plex CosMx spatial transcriptomics analysis on an independent cohort of 8 EBV-positive and 12 EBV-negative DLBCL patient samples (**Fig. 5E, top row**). We observed heterogeneous spatial organization between tumor cells, macrophages, and T cells, with T cells in the EBV-positive TME consistently exhibiting elevated dysfunction signatures compared to their EBV-negative counterparts (**Fig. 5E, bottom** and **Supp Fig. 7E**), confirming our findings from the primary IN-DEPTH cohort.

Coupling IN-DEPTH with SGCC extended our proposed mechanism (**Fig. 4K**) to reveal two distinct spatially orchestrated cellular circuits in the DLBCL TME: In EBV-positive cases, tumor cells preferentially associate with macrophages to condition an immunosuppressive environment that impairs CD4 T cell function. Conversely, in EBV-negative TMEs, increased macrophage-CD4 T cell interactions foster a pro-inflammatory TME (**Fig. 5F**).

## Discussion

IN-DEPTH addresses current limitations in spatial multiomics efforts by enabling same-slide protein and RNA profiling to significantly expand the number of simultaneously detectable biomolecules without proportional increases in cost and time. This approach eliminates the need for challenging computational integration of adjacent tissue slides and associated artifacts (**Supp Fig. 1A**). Our spatial protein-first strategy enables targeted spatial transcriptomics dissection guided by biological context, offering a resource-effective alternative to whole-slide transcriptome profiling in a platform agnostic manner. While IN-DEPTH accommodates various commercially available or home-brewed spatial platforms, careful consideration is essential. For instance, the tyramide signal amplification approach in the Polaris involves direct covalent deposition of Opal fluorophores on the tissue (24), and may require significant photobleaching or alternatives for compatibility with fluorescence in-situ hybridization-based spatial transcriptomics platforms such as the CosMx. Importantly, IN-DEPTH is carefully optimized to maximize tissue integrity while enabling robust protein epitope staining via spatial proteomics, RNA signal retention, and subsequent H&E staining for downstream pathological verification (**Figs. 1C, 2B, 4B** and **Supp Fig. 2C**), and thus potentially allowing for additional spatial modalities on the same slide beyond protein and RNA (55).

SGCC is derived through graph signal processing and GFT-based mathematical reasoning (**Fig. 3**) and serves as a measure for quantifying the relative spatial positions of any two cell phenotypes in the low-frequency domain. Due to the unbiased and interpretable nature of GFT, this metric can effectively gauge spatial relationships between two cell phenotype distributions. Furthermore, when multiple samples or multiple ROIs are available, SGCC can be treated as a continuous or ordinal spatial factor. In this capacity, it can be integrated with transcriptomic data to identify covarying genes, thus offering a comprehensive understanding of the relationship between cellular arrangements and functional states.

We demonstrate the utility of IN-DEPTH in dissecting EBV-associated immune modulation in the DLBCL TME, revealing key contrasting features including tumor-associated increase in M2-like macrophages with diminished HLA-DR and increased PD-L1 expression linked to increased CD4 T cell dysfunction in EBV-positive DLBCL (**Fig. 4**). While this biological mechanism has not been described in EBV-positive DLBCL, it is supported by data from prior studies across various biological systems (56–61). Our findings additionally support the prevalent and consistent association between EBV positivity, poor prognosis, and inferior outcomes in DLBCL (62–69), as well as the relationship between immunosuppressive macrophages and ineffective immune responses in other cancers (30, 70– 76). We apply IN-DEPTH here to contextualize the functional diversity of macrophages *in situ*, which coupled with SGCC integrative analysis uniquely enabled additional functional assessment of macrophages based on their spatial organization within the tumor-macrophage-CD4 T cell immunomodulatory axis, uncovering a tumor virus-dependent rewiring of this tripartite interaction in the DLBCL TME (**Fig. 5**). The differences in EBV-stratified T cell immune dysfunction may in part explain the different responses to immune checkpoint blockade in DLBCL (57, 77, 78). The ability of IN-DEPTH and SGCC to decipher nuanced cellular functional states demonstrated here highlights their potential in advancing our understanding of spatially organized immune interactions and their impact on tumor progression and immune dysfunction.

Several exciting opportunities exist for further development. The preservation of tissue integrity enables future integration of histological features and other spatial modalities. We focused here on cell-type specific transcriptome capture for resource efficiency, but expansion to single-cell or subcellular resolution is certainly possible (**Fig. 1C**) and will necessitate additional advancements in computational approaches. IN-DEPTH datasets will also be fundamental as ground truth in future computational developments for a variety of tasks, including bulk deconvolution, multi-modal integration, and beyond. The experimental and computational advances presented herein demonstrate the potential for comprehensive tissue analysis with new insights gained through same-slide integrated spatial multi-omics. We anticipate this approach to be broadly applicable across spatial platforms, to accelerate discovery and mechanistic research across multiple diseases.

## Materials & Methods

### Human Tissue Acquisition and Patient Consent

All formalin-fixed paraffin-embedded (FFPE) tissues used in this study were sectioned 5 µm thick on SuperFrost glass slides (VWR, 48311-703) and obtained from the following sources. The tonsil tissues in **Figs. 1 & 2** were generously provided by S.J.R. from the Brigham and Women’s hospital (IRB# 2016P002769 and 2014P001026), the DL-BCL tissue for SignalStar-GeoMx (**Fig. 1C, row 3**) was purchased from amsBio (amsBio, AMS-31010), the kidney cancer (**Fig. 1C, row 4**) and lymph node tissues (**Fig. 1C, row 5**) were generously provided by S.S. from the Dana Farber Cancer Institute (IRB# DFCI 13-425), the periodontal disease tissue for CODEX-VisiumHD (**Fig. 1C, row 6**) was generously provided by D.M.K. from Harvard Dental School (IRB# 22-0587), the DLBCL tissue for CODEX-CosMx (**Fig. 1C, row 8**) was obtained from W.R.B. from University of Rochester Medical Center (IRB# STUDY159), and the uterine cancer tissues (**Supp Fig. 1B**) were generously provided by B.H. from Stanford University Medical School.

For comparing EBV-positive vs EBV-negative DLBCL (**Fig. 4**), 30 patient samples (17 EBV-positive, 13 EBV-negative) were sectioned from two tissue microarrays (TMA). The Dana-Farber Cancer Institute TMA, constructed by S.S. and S.J.R. (IRB# 2016P002769 and 2014P001026), includes 1 core from each patient (10 EBV-positive, 9 EBV-negative) and 1 tonsil control core, with each core measuring 1.5 mm in diameter. The University of Rochester Medical Center TMA, constructed by D.N., P.R., and W.R.B. (IRB# STUDY159), includes 1 core from each patient (13 EBV-positive, 6 EBV-negative) and 1 tonsil control core, with each core measuring 2.0 mm in diameter. For validating EBV-positive vs EBV-negative signatures (**Fig. 5E**), one 1.5mm diameter core from each of 18 patient samples (8 EBV-positive, 10 EBV-negative) were sectioned from a TMA from University Hospital and Comprehensive Cancer Center Tübingen that was constructed by L.F., L.K., and C.M.S. EBV status for all DLBCL biopsies were verified using in-situ hybridization for EBER as part of the routine clinical pathology process. Detailed de-identified information for the DLBCL patients are in **Supp Table 5**.

### Antibody Panel Selection, Conjugation, and Titration

Antibodies used in the CODEX experiments were conjugated in-house and include previously validated antibody clones (10, 21, 33). In brief, the specificity of antibody candidates were first validated via immunohistochemistry (IHC) on FFPE cell pellets or FFPE lymphoid tissues to ensure robustness of staining. The selected antibody clones were then conjugated by either maleimide, lysine, or biotinylation chemistries, and each conjugated antibody was titrated and validated via immunofluorescence on FFPE lymphoid tissues. Readers of interest are referred to the following publications for a more detailed guide on antibody target selection and optimization (20, 79). Antibodies used for the SignalStar, Polaris, and Orion experiments were obtained from their respective commercial sources. Details regarding the antibody clones, vendors, conjugated channels, titers, exposure times, and assigned channels throughout the study are in **Supp Table 6**.

Maleimide-based conjugations were performed with minor modifications from a previously published protocol (28). Briefly, 50 or 100 µg of carrier-free antibody was concentrated using a PBS-T pre-wetted 50kDa filter (Sigma Millipore, UFC5050BK) and then incubated with 0.9 µM TCEP (Sigma, C4706-10G) for 10-30 minutes in a 37°C water bath to reduce the thiol groups for conjugation. Reduction was quenched by two washes with Buffer C (1mM Tris pH 7.5, 1mM Tris pH 7.0, 150mM NaCl, 1mM EDTA) supplemented with 0.02% NaN3. Maleimide oligos were resuspended in Buffer C supplemented with NaCl (Buffer C, 250mM NaCl). The reduced antibody was next incubated with 100 or 200 µg (for 50 or 100 µg of antibody, respectively) of maleimide oligos (Biomers, 5’-Maleimide) in a 37°C water bath for 2 hrs. The resulting conjugated antibody was purified by washing for three to five times with the 50kDa filter with high-salt PBS (1× DPBS, 0.9M NaCl, 0.02% NaN3). The conjugated antibody was quantified in IgG mode at A280 using a NanoDrop (Thermo Scientific, ND-2000). The final concentration was adjusted by adding >30% v/v Candor Antibody Stabilizer (FisherScientific, NC0414486) supplemented with 0.2% NaN3, and the antibody was stored at 4°C.

Lysine-based conjugations were performed with minor modifications from the official Alexa Fluor™ 532 / 594 / 647 Labeling Kit protocols (ThermoFisher, A20182 & A20185 & A20186). Briefly, 100 µg of carrier-free antibody was adjusted to a concentration of 1 mg/mL and mixed with 10 µL of 1M sodium bicarbonate buffer with gentle agitation for 5 min. The basic pH antibody was then transferred into the Alexa Fluor™ reactive dye with gentle pipetting to dissolve the dye. The labeling reaction proceeded in the dark for 1 hr at room temperature (RT), and the vial was gently inverted 5 times every 15 min. A purification resin bed was prepared by thoroughly resuspending the resin by violent agitation, and then centrifuging the resin through the provided filters at 1200 ×g for 8 min until there was minimal residual buffer remaining in the resin bed. The conjugated antibody was then pipetted into the resin bed and allowed to absorb into the bed for 1 min. The antibody was collected by centrifuging at 1200 ×g for 5 min and then stored at 4°C.

Biotinylation was performed using a commercial rapid biotinylation kit (Biotium, 92244) according to manufacturer’s instructions. Briefly, 75 µg of carrier-free antibody was biotinylated, with a conjugation time of 15 min. The conjugated antibody was diluted in 300 µL provided Storage Buffer and then stored at 4°C.

### Spatial Proteomics: Antibody Staining and Imaging

The tissue antigen retrieval and photobleaching steps were standardized across all spatial proteomics assays accordingly. Briefly, FFPE tissue slides were baked in an oven (VWR, 10055-006) at 70°C for 1 hr, then thoroughly deparaffinized by immersing in xylenes for 2× 5 minutes. The slides were then subject to a series of graded solutions for rehydration using a linear stainer (Leica Biosystems, ST4020), with each step proceeding for 3 min: 3× xylene, 2× 100% EtOH, 2× 95% EtOH, 1× 80% EtOH, 1× 70% EtOH, 3× UltraPure water (Invitrogen 10977-023), and finally left in UltraPure water (Invitrogen 10977-023). Antigen retrieval was then performed at 97°C for 20 min with pH 9 Target Retrieval Solution (Agilent, S236784-2) using a PT Module (ThermoFisher, A80400012), after which the slides were cooled to room temperature on the benchtop and washed in 1× PBS for 5 min. Tissue regions were circled with a hydrophobic barrier pen (Vector Laboratories, H-4000), rinsed in 1× PBS to remove residual ink, then washed in 1× TBS-T prior to photobleaching and antibody blocking. For assays that include staining with a biotinylated antibody, an extra biotin blocking step was included at this point with a commercial Biotin Blocking kit (Biolegend, 927301). Briefly, slides were first incubated with the avidin solution for 30 min at RT followed by two quick rinses 1× TBS-T and one 2 min wash with 1× TBS-T, and next incubated with the biotin solution for 30 min at RT followed by two quick rinses 1× TBS-T and one 2 min wash with 1× TBS-T. Photobleaching and antibody blocking was then performed by first washing the slides in S2 Buffer (2.5 mM EDTA, 0.5× DPBS, 0.25% BSA, 0.02% NaN3, 250 mM NaCl, 61 mM Na2HPO4, 39 mM NaH2PO4) for 20 min, then blocking using BBDG (5% normal donkey serum, 0.05% NaN3 in 1× TBS-T wash buffer (Sigma, 935B-09)) supplemented with 50 µg/mL mouse IgG (diluted from 1 mg/mL stock (Sigma, I5381-10mg) in S2), 50 µg/mL rat IgG (diluted from 1 mg/mL stock (Sigma, I4141-10mg) in S2), 500 µg/mL sheared salmon sperm DNA (ThermoFisher, AM9680), and 50 nM oligo block (diluted from stock with 500 nM of each oligo in 1× TE pH 8.0 (Invitrogen, AM9849). The blocking occurred in a humidity chamber on ice while being photobleached for 90 min using Happy Lights (Verilux, VT22), with the temperature continuously monitored to ensure that it was kept below 40°C. After photobleaching and antibody blocking, tissues were stained and imaged accordingly based on the respective assay, as described below. Note that the photobleaching and blocking setup was different for the Orion (more details below).

#### CODEX

Tissues were stained for 1 hr at RT in a humidity chamber, and then washed in S2 Buffer twice for 2 min each at RT. The slides were first fixed in 1.6% PFA (diluted from 16% stock (EMS Diasum, 15740-04) in S4 Buffer (4.5 mM EDTA, 0.9× DPBS, 0.45% BSA, 0.02% NaN3, 500 mM NaCl)) twice for 5 min each at RT, after which the slides were rinsed twice in 1× PBS followed by a 2 min wash in 1× PBS. The slides were next fixed with ice-cold methanol for 5 min on ice (while intermittently lifted to scrape off the hydrophobic barrier using a cotton-tipped applicator starting from the 3 min timepoint), after which the slides immediately rinsed twice in 1× PBS followed by a 2 min wash in 1× PBS. The slides were finally fixed in 4 µg/µL of BS3 Final Fixative (diluted from 200 µg/µL stock (ThermoFisher, 21580) in 1× PBS) twice for 10 min each in the dark at RT, after which the slides were rinsed twice in 1× PBS followed by a 2 min wash in 1× PBS.

To prepare the slides for imaging in the automated PhenoCycler Fusion platform (**Fig. 1C, row 1**), flow cells (Akoya Bioscience, 240205) were mounted by securely pressing them on each tissue slide for 30 s, followed by 10 min of incubation in 1X CODEX Buffer. A reporter plate was also prepared for each tissue slide such that each well corresponds to each imaging cycle. Briefly, a 96-well black reporter plate (BRAND Tech, 781607) was prepared by filling each well with plate buffer (500 µg/mL sheared salmon sperm DNA in 1× CODEX buffer (10mM Tris pH 7.5, 0.02% NaN3, 0.1% Triton X-100, 10 mM MgCl2-6H2O, 150mM NaCl)) supplemented with 1:300 (54.11 mM) of Hoechst 33342 (ThermoFisher, H3570), and adding complementary reporter oligos conjugated with ATTO550 or AlexaFluor647 (GenScript, HPLC purified) to a final concentration of 100 nM each. The wells were then sealed using aluminum plate seal (ThermoFisher, AB0626) and mixed by inverting the plate several times. Low DMSO (80% 1× CODEX buffer, 20% DMSO) and High DMSO (10% 1× CODEX buffer, 90% DMSO) buffers were also prepared fresh each run by mixing 1× CODEX Buffer in DMSO (Sigma, 472301-4L), which was used by the PhenoCycler Fusion to strip and hybridize the reporter oligos. After imaging, the flow cell was removed prior to RNA probe hybridization by using a razor blade to pry the flow cell and gently scrape off any adhesive while repeatedly dipping in 1× PBS. Personal protective equipment was worn at all times at this step. After the flow cell and adhesive were removed, slides were washed twice in 1× PBS.

For the data acquired by manual cycling imaging (**Fig. 1C, row 2**), the slides were first rinsed in 1× CODEX Buffer followed by an initial stripping cycle in stripping buffer (25% 10x CODEX Buffer, 75% DMSO) twice for 5 min each. The slides were subsequently washed twice in 1× CODEX buffer for 5 min each, incubated for 10 min with plate buffer supplemented with 100 nM SYTO13 (ThermoFisher, S7575), then washed twice again for 5 min each in 1× CODEX buffer. The slides were then loaded into the GeoMx and scanned as the initial blank cycle. Subsequent cycles were carried out as follows: 2× 5 min incubation in stripping buffer, washing twice in 1× CODEX for 5 min each, 10 min incubation in plate buffer supplemented with 100nM SYTO13 and three 100nM reporter oligos conjugated to Alexa Fluor 532, 594, or 647 (GenScript, HPLC purified), and finally washing in 1× CODEX Buffer twice for 5 min each. After all marker cycles, a final blank cycle stained with only 100 nM SYTO13 was also included to ensure clearance of signal. All steps were performed at RT on the benchtop, all stripping and washing steps were performed in polypropylene Coplin jars (Tedpella, 21038), while all reporter oligo incubations were performed in a humidity chamber. For all imaging, slides were loaded into the provided slide holder in the GeoMx and hydrated with 3 mL of Buffer S prior to operating the instrument. After imaging, slides were washed twice in 1× PBS.

#### SignalStar

The SignalStar reaction occurs in two rounds with four antibodies imaged per round, and was performed using the commercial buffers (Cell Signaling Technology, 63043S) unless otherwise mentioned. Briefly, during each round, tissues were first incubated with SignalStar Amplification Solution 1 (1:100 of each SignalStar complementary oligo diluted in amplification buffer) for 2 hr (round 1 that includes 1:100 of each antibody) or 40 min (round 2 that does not contain antibodies) at 4°C, and then rinsed in 1× TBS-T for 30 s. Tissues were then fixed in 4% PFA (diluted from 16% stock (EMS Diasum, 15740-04) in 1× PBS) for 5 min at RT. After washing using UltraPure water (Invitrogen 10977-023), eight rounds of amplification was performed accordingly using the corresponding amplification solution (1:50 of each amplification oligo diluted in amplification buffer), with a 30 s UltraPure water (Invitrogen 10977-023) rinse between each round of amplification. A 20 min ligation step was performed accordingly using SignalStar Ligation Solution (50% Ligation Buffer, 2% T4 ligase (from a stock “5 units per mL”), and 1 mM ATP prepared using UltraPure water (Invitrogen 10977-023)), followed by another 30 s Ultrapure water (Invitrogen 10977-023) rinse. Tissues were then stained with 1:300 of Hoechst 33342 (ThermoFisher, H3570) for 5 min at RT, rinsed with 1× TBS-T, and coverslipped with ProLong™ Gold Antifade Mountant (P36930). Tissues were then imaged on the corresponding 4-color channels using the PhenoCycler Fusion platform. After imaging, the coverslip was removed by dipping in 1× TBS-T followed by incubation with the SignalStar Fluorescent Removal Solution for 2 hr at 37°C and rinsed with UltraPure water (Invitrogen 10977-023) for 30s. To ensure complete removal of signal, tissues were stained with 1:300 of Hoechst 33342 (ThermoFisher, H3570) for 5 min at RT and then imaged again. The coverslip was similarly removed by dipping in 1× TBS-T. After both SignalStar reactions, slides were finally washed five times in 1× PBS to ensure complete removal of glycerol.

#### Polaris

An optimized tissue staining assay was performed on a Bond RX Autostainer (Leica Biosystems) using the Akoya Biosciences Opal tyramide signal system. The antibody:fluorophore pairings are: CD8 on Opal Polaris 480 (1:50), PD-1 on Opal Polaris 690 (1:100), TIM-3 on Opal Polaris 620 (1:150), LAG-3 on Opal Polaris 570 (1:50), CD20 on Opal Polaris 520 (1:150), and CD163 on Opal Polaris 780 (1:25)/TSA-DIG (1:100). Prior to imaging, slides were mounted using 1× PBS and sealed with nail polish. Whole-slide multispectral images were acquired at 20× magnification using the PhenoImager HT automated quantitative pathology imaging system (Akoya Biosciences), while implementing the Inform 3.0 software was then used to deconvolute the multispectral images. After imaging, a cotton swab dipped with xylenes was used to remove the nail polish and unmount the coverslip, and slides were then washed twice in 1× PBS.

#### Orion

After antigen retrieval, the autofluorescence quenching, blocking, and antibody staining steps were instead performed according to the manufacturer’s protocol. After antibody staining, tissues were coverslipped using 1× PBS and sealed with nail polish. Whole-slide images were acquired using the Orion (Rarecyte). After imaging, a cotton swab dipped with xylenes was similarly used to unmount the tissue, followed by washing twice in 1× PBS.

### Spatial Proteomics: Cell Segmentation and Annotation

The following paragraphs describe real-time analyses of the multiplexed images that were performed in parallel with the overnight RNA probe hybridization after image acquisition. Note that these steps are only performed for **Fig. 2** and **Fig. 4**. Details of the thorough analyses performed after completing the IN-DEPTH experiment are described in the Spatial Proteomics Analysis section.

#### Cell segmentation

For both the tonsil (**Fig. 2**) and DL-BCL (**Fig. 4**) datasets, cell segmentation was only performed on the CODEX image using the MESMER model of DeepCell (v0.12.2) (80, 81), with maxima_threshold set to 0.075 and interior_threshold set to 0.05. The nuclear channel input of MESMER was DAPI for both datasets. The membrane channel input of MESMER for the tonsil dataset (**Fig. 2**) was a summation of CD11b, CD68, CD20, CD163, CD31, and CD3, while for the EBV-positive vs EBV-negative DLBCL dataset (**Fig. 4**), it was a summation of HLA1, HLA-DR, and CD31.

#### Image registration between CODEX and GeoMx

Scale-Invariant Feature Transform (SIFT) algorithm was used (82) for feature detection and feature description of the Fusion DAPI image and the GeoMX SYTO13 image. Then, a brute-force matcher was used to match the features between the two images. A ratio test was used to determine if a specific match should be considered as a “good match”. The source point (the CODEX image) and the destination point (the GeoMx image) of the “good matches” were used to calculate the affine transformation matrix that would register the CODEX image’s coordinates into the GeoMx image’s coordinate system. The software used and the specific hyperparameters for the algorithm and ratio test are in **Supp Table 7**.

#### Single-cell feature extraction

For each marker, the pixel value within the area of each cell (determined by the segmentation mask) was summed and then divided by the area of each cell, and the resulting cell-size scaled sum was set as the expression value for a given marker. For the DLBCL dataset (**Fig. 4**) where 3 markers were acquired on the GeoMx, the segmentation mask generated from the CODEX image was applied to the GeoMx image to ensure that the same cell imaged between the two instruments contained the same cell label, from which the cell features were similarly extracted and scaled to cell size. Finally, the scaled single-cell features extracted from the Fusion and GeoMx images were joined together by cell label and tissue core ID.

#### Cell phenotyping

The extracted features were first scaled to a standardized range of [0,1], and cell phenotyping was then performed through an iterative clustering and annotating process with PhenoGraph (83). For the tonsil dataset (**Fig. 2**), the 12 phenotyping markers used were CD20, Pax5, BCL6, CD3, CD8, CD4, FoxP3, CD11c, CD31, CD68, CD163, and CD11b, which allowed the annotation of BCL6+ B cells, BCL6-B cells, CD4 T cells, CD8 T cells, endothelial cells, Tregs, dendritic cells (DCs), M1-like macrophages, M2-like macrophages, and other myeloids. For the EBV-positive vs EBV-negative DLBCL dataset (**Fig. 4**), the phenotyping markers used were CD20, Pax5, CD3, CD8, CD4, FoxP3, CD11c, CD31, CD68, and CD163, which allowed the annotation of CD4 T cells, CD8 T cells, endothelial cells, Tregs, DCs, M1-like macrophages, M2-like macrophages, and tumor cells. Cells that showed unclear marker enrichment patterns were annotated as “Other” cells.

During the annotation process, clustering results were first visualized using a heatmap showing the Z-score of each marker within each cluster. This was used as a basis to annotate each cluster based on their marker Z-score combinations while visually inspecting the original images to confirm annotation accuracy. After an initial round of clustering with PhenoGraph was performed, clusters with clear enrichment patterns were annotated, while clusters with mixed patterns underwent additional rounds of clustering and annotation using a targeted set of phenotyping markers. This process was iterated until all identifiable cells were annotated. To visualize and confirm the assigned annotations, Mantis Viewer (84) was utilized to overlay the annotation onto the segmentation mask and the marker image for visual inspection. The final annotations were then examined by visually inspecting with multiplexed images and H&E stains and verified by S.K. and S.J.R..

For the Tonsil experiment (**Fig. 2**), we annotated one tissue section using the above-described procedure. Leveraging upon the advantage of adjacent tissue sections and the reproducible high-quality tissue staining, annotation of the the adjacent section was guided by MAPS (85), followed by further refinement using the same procedures as described above.

### Spatial Transcriptomics: Probe Hybridization and Transcriptome Capture

At this point, all tissues were equilibrated in 1× PBS, including the control slides that were paused after antigen retrieval. Tissues were then hybridized for transcriptome capture accordingly based on the respective assay, as described below.

#### GeoMx

The RNA probe staining cocktail was prepared using the Nanostring RNA Slide Prep kit (Nanostring, 121300313) using the Nanostring Human Whole Transcriptome Atlas detection probe set (Nanostring, 121401102). The RNA probe cocktail was then applied to the tissue slides, sealed with a hybridization cover slip (EMS Diasum, 70329-40), and incubated overnight (around 18 hrs) at 37°C. After RNA probe hybridization, tissue slides were first washed twice in Stringent Wash Buffer (2× saline-sodium citrate (SSC) (Millipore Sigma, S6639) in 50% formamide (Millipore Sigma, 344206-1L-M) for 5 min each at 37°C, and subsequently washed twice with 2× SSC for 5 min each at RT on a belly dancer. Tissues were then stained with SYTO13 (100 nM) for 10 min at RT, and washed twice in 2× SSC for 2 min each at RT to visualize nuclear morphology. Slides were then scanned on the GeoMx for region of interest (ROI) selection, while ensuring that the IN-DEPTH stained and control slides were always scanned in parallel. Square 484×484 µm ROIs were drawn for each experiment: 18 in **Fig. 1C rows 1-2**, 24 in **Fig. 1C row 3**, 16 in **Fig. 1C row 4**, 8 in **Fig. 1C row 5**, and 25 in **Supp Fig. 1B**.

For the tonsil biological validation component (**Fig. 2**), a few adjustments were incorporated. Sixteen 660×760 µm rectangular ROIs were selected on each adjacent tissue section with emphasis on lymphoid nodules (**Fig. 2B** and **Supp Fig. 2B**). The location of each ROI on the GeoMx was then recorded by their four vertices, and these coordinates were used to crop out one sub-region for each ROI from the CODEX-to-GeoMx registered full-tissue segmentation mask. Within each sub-region for each ROI, a segmentation mask for each annotated cell population was iteratively generated to enable cell-type specific RNA collection. Each cell-type specific segmentation mask was then converted into a binary mask by setting the pixel value of all the cell areas to 255 and pixel value for all background areas to 0. These masks were then re-uploaded onto the GeoMx instrument to guide cell-type specific RNA genome-wide transcriptome extraction, ranked from the lowest to highest cell proportion within each ROI, such that transcript collection would proceed in this order.

For the EBV-positive vs. EBV-negative DLBCL component (**Fig. 4**), more adjustments were incorporated. The Nanostring Human Whole Transcriptome Atlas detection probe was combined with a custom spike-in panel of probes against 14 targeted EBV genes (*EBER1, EBER2, EBNA1, EBNA2, EBNALP, LMP1, RPMS1, BALF1 BCRF1, BHRF1, BNLF2A, BNLF2B, BNRF1, BZLF1*). After 2× SSC and formamide washing, slides were stained with antibodies against Tox1/2, c-Myc for 1 hr at RT, followed by SYTO13 (100 nM) streptavidin (used to visualize the biotinylated PD-L1 antibody) for 10 min at RT. The stained slides were then washed twice in 2× SSC for 2 min each at RT prior to GeoMx scanning. One 660×785 µm rectangular ROI was drawn for each patient core with emphasis on tumor-enriched regions. The location of each ROI on the GeoMx was similarly recorded by their four vertices and used to crop out the corresponding sub-regions, from binary 0/255 segmentation masks for each annotated cell population were iteratively generated, ranked, and uploaded onto the GeoMx for transcriptome extraction.

After transcriptome capture, unique molecular barcodes for the RNA probes were aspirated from each cell population to 96-well collection plates (Nanostring, 100473), except for the first aspirate for each plate which is the default negative control. Collection plates that were fully filled were dried according to official Nanostring protocol and stored at -20°C until transcript collection for all other collection plates within each experiment was completed. Sequencing library preparation was then performed starting from the dried collection plates. Each aspirate was first resuspended in 10 µL of UltraPure water (Invitrogen 10977-023) and then uniquely indexed using the Illumina i5×i7 dual indexing system as part of the Nanostring NGS library preparation kits (Nanostring, 121400201 & 121400202 & 121400203 & 121400204). The PCR reaction was prepared in 96-well PCR plates (ThermoFisher 4306737), where each well contained 4 µL of aspirate, 1 µM of each i5 and i7 primers, and 1× library preparation PCR Master Mix, adding up to 10 µL per well. The PCR reaction conditions were 37 °C for 30 min, 50 °C for 10 min, 95 °C for 3 min, followed by 18 cycles of 95 °C for 15 s, 65 °C for 60 s, 68 °C for 30 s, followed by a final extension of 68 °C for 5 min before holding indefinitely at 12°C. Next, 4 µL of PCR product from each well was pooled into DNA LoBind tubes (Eppendorf 022431021) for purification, with 1 LoBind tube used per collection plate. For the first round of purification, 1.2× volume of AMPure XP beads (Beckman Coulter A63881) were first added to the pooled PCR products and incubated at RT for 5 min. Beads were then pelleted on a magnetic stand (ThermoFisher 12321D), washed twice with 1 mL of 80% ethanol, and eluted with 54 µL of elution buffer (10 mM pH 8.0 Tris-HCl, 0.05% Tween-20). The second round of purification was performed using 50 µL of eluted DNA from the first round, incubated with 1.2× volume of AMPure XP beads and washed twice in 1 mL of 80% ethanol. A final elution was done at 2:1 ratio of aspirate (number of wells) to elution buffer (volume in µL), and 0.5 µL of the final eluate was diluted in 4.5 µL of UltraPure water (Invitrogen 10977-023) (1:10 dilution) to confirm library purity and concentration on the Agilent TapeStation.

For each experiment, the same concentration of each sub-library (eluted in individual DNA LoBind tubes) was pooled into one LoBind tube to be sent for next-generation sequencing. PhiX sequencing control (Illumina FC-110-3002) was added into the library, with amount adjusted based on the percentage of total reads allocated for PhiX as per the sequencing platform used (5% on the NovaSeq X Plus, 20% on the NextSeq2000). Paired-end sequencing was then performed on the NovaSeq X Plus (Tonsil tissue experiments, **Figs. 1 & 2**) or NextSeq2000 (DLBCL experiment, **Fig. 4**), with a total sequencing depth calculated as:

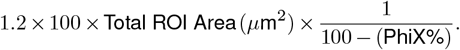

#### VisiumHD

Slides were first subjected to H&E staining and imaging as described in the next section. Afterwards, tissues were dried at 37°C for 3 min using a thermal cycler. Tissues were then destained with 0.1 M HCl at 42°C for 15 min, followed by 3× washes and incubations with TE buffer, and finally submerged in 1× PBS.

As the default VisiumHD workflow has a de-crosslinking step prior to probe hybridization, the control VisiumHD-only slide was subjected to de-crosslinking at 80°C for 30 min using the Decrosslinking Mix provided by the manufacturer followed by probe hybridization at 50°C overnight following manufacturer protocols (10X Genomics #1000668 and #1000466). For the CODEX-VisiumHD slide, tissues were incubated with 2 µg/mL Proteinase K (Thermo Fisher Scientific, AM2546) prepared with 1× PBS at 40C for 20 min, followed by three washes in UltraPure water (Invitrogen 10977-023). Tissues were then fixed in 10% NBF (EMS Diasum, 15740-04) at RT for 1 min, and the fixation process was stopped by incubating the tissue twice in NBF stop buffer (0.1M Tris and 0.1M Glycine) for 5 min each at RT, followed by a 1× PBS wash for 5 min at RT. The tissues were then similarly subjected to probe hybridization (10X Genomics #1000466) at 50°C overnight following manufacturer protocols.

Following post-hybridization wash, the tissues were subjected to probe ligation at 37°C for 1 hr,washed with post-ligation wash (10X Genomics #1000668) at 57°C for 5 min, and finally with 2× SSC buffer. The tissues were then stained with 10% Eosin at RT for 1 min and washed with 1× PBS. The tissues were loaded into the Visium CytAssist, adjusted to align with the slide subjected to Visium HD, followed by probe release. Two square 6.5×6.5 mm ROIs were drawn for this experiment in **Fig. 1C, row 6** due to the inherent size of each cassette (10X Genomics #1000669 and #1000670). Probes were then extended with a thermal cycler and eluted with 0.08 M KOH. Probes from each of the tissue samples were amplified with individual Dual Index TS Set A (10X Genomics #PN-1000251) in a thermal cycler followed by PCR-clean up with SPRIselect Reagent (Beckman Coulter #B23317). The libraries were QC-ed through High Sensitivity DNA Assay (Agilent Technologies) and sequenced paired-end on a HiSeq2000 (Illumina).

#### CosMx

An incubation frame was first applied on each slide to ensure that liquid remains on the tissue surface. Tissues were then digested with 2 µg/mL Proteinase K (Thermo Fisher Scientific, AM2546) prepared with 1× PBS for 20 min at 40°C, followed by three washes in UltraPure water (Invitrogen 10977-023). Fiducial solution (0.001% of fiducials in 2× SSC-T) was applied afterwards for 5 min at RT, which is immediately followed by tissue fixation in 10% NBF (EMS Diasum, 15740-04) for 1 min at RT. The fixation process was quenched twice in NBF stop buffer (0.1M Tris and 0.1M Glycine) for 5 min each at RT, followed by a 1× PBS wash for 5 min at RT. To block nonspecific probe and antibody binding, a 100 mM NHS-acetate mixture was prepared immediately prior to application and incubated for 15 min at RT in a humidified chamber. Slides were then washed twice in 2× SSC for 5 min each at RT.

The RNA detection probes were prepared by denaturing at 95°C for 2 min using a preheated thermal cycler and then immediately chilled in an ice bucket for 1 min. Note that different detection probe panels were used, with a 1k panel for **Fig. 1C, row 7** and a 6k panel for **Fig. 1C row 8**. Afterwards, the RNA probe cocktail was prepared according to manufacturer guidelines. The upper layer of the incubation frame was carefully removed to apply the probe cocktail while ensuring the liquid remains within the incubation frame boundary without any bubbles introduced, after which an incubation frame cover was used to seal the RNA probe cocktail within. Probes were allowed to hybridize at 37°C for 16 hrs. After RNA probe hybridization, tissue slides were first washed twice in Stringent Wash Buffer (2× saline-sodium citrate (SSC) (Millipore Sigma, S6639) in 50% formamide (Millipore Sigma, 344206-1L-M)) for 25 min each at 37°C, and subsequently washed twice with 2× SSC for 5 min each at RT on a belly dancer. Tissues were then stained with SYTO13 (100 nM) buffered in blocking buffer for 15 min at RT, washed in 1× PBS for 5 min, followed by staining with a designated antibody cocktail for 1 hr at RT to demarcate cell boundaries. After antibody staining, slides were washed thrice in 1× PBS followed by another round of incubation using freshly-prepared NHS-acetate mixture for 15 min at RT. Slides were then washed twice in 2× SSC for 5 min each at RT. Slides were then scanned on the CosMx for region of interest (ROI) selection, while ensuring that the IN-DEPTH stained and control slides were always scanned in parallel. Square 500×500 µm ROIs were drawn for each experiment: 36 in **Fig. 1C, row 7**, and 18 in **Fig. 1C, row 8**.

### Hematoxylin & Eosin Staining and Imaging

#### VisiumHD

H&E staining was part of the VisiumHD protocol. Slides were first immersed twice in UltraPure water (Invitrogen 10977-023) for 20 s each. H&E staining was performed a serial incubation in hematoxylin (StatLab, HXMMHPT), blueing buffer (StatLab HXB00588E), and eosin (StatLab STE0243) for 1 min each at RT, with three UltraPure water (Invitrogen 10977-023) washes between each incubation. Next, glycerol was used to coverslip the VisiumHD only slide while UltraPure water (Invitrogen 10977-023) was used to coverslip the Codex-VisiumHD slide. Slides were then scanned using the Grundium Ocus40 slidescanner (Grundium MGU-00003). After scanning, the coverslip was removed by immersing the slides in UltraPure water (Invitrogen 10977-023) and continued with drying and destaining and detailed in the previous section.

#### GeoMx & CosMx

All slides were stored in 2× SSC at 4°C after transcriptome capture for H&E staining to visualize and confirm tissue morphology immediately after completing quality control evaluation of the captured transcripts. Slides were first equilibrated in UltraPure water (Invitrogen 10977-023) at RT prior to staining with Modified Mayer’s Haematoxylin (StatLab HXMMHPT) for 5 min at RT, followed by rinsing thrice with UltraPure water (Invitrogen 10977-023). Slides were then treated with Bluing Solution (StatLab HXB00588E) to develop the blue coloration, and subsequently rinsed thrice with UltraPure water (Invitrogen 10977-023) at RT. The slides were then equilibrated in 95% ethanol for 1 min prior to staining with a solution of Eosin Y and Phloxine B (StatLab STE0243) for 1 min, followed by rinsing by dipping 12 times each in three changes of fresh 95% ethanol. Finally, the slides underwent graded dehydration by dipping once in 70% ethanol, once in 100% ethanol, and once in two changes of xylenes. Excess xylenes was gently dabbed off and glass coverslips (Creative Waste Solutions CSM-2450) were mounted with xylene-based mounting medium (OptiClear Xylene, SSN Solutions, CSM1112). The slides were left to dry overnight at RT, after which they were scanned using the Grundium Ocus40 slidescanner (Grundium MGU-00003). The H&E stains were verified by S.K. and S.J.R. for tissue quality and morphological consistency with the multiplexed spatial proteomics images.

### Spatial Transcriptomics: Batch Correction

#### GeoMx data

The demultiplexed FASTQ output files from next-generation sequencing were used to map and quantify the human probes (and EBV probes for DLBCL data) through the GeoMx Data Analysis software pipeline (8). The .dcc files produced were then uploaded onto the GeoMx to generate gene counts tables using the default “QC” and “Biological probe QC” settings without filtering out any genes.

The original cell-type annotations distinguished multiple T cells (CD4 memory, CD4 naive, CD8 memory, CD8 naive), macrophage (M1-like, M2-like), endothelial, and several tumor subtypes (including subsets defined by BCL2, BCL6, and Myc expression level), as shown in **Supp Fig. 5**. To streamline the analyses, closely related cell subsets were merged into broader categories: memory and naive T cell subpopulations were combined into respective CD4 or CD8 T cells, and tumor subpopulations (originally BCL2+, BCL6+, Myc+, and other tumors) were aggregated to represent a collective malignant B-cell population. Following the merging of related cell subpopulations, gene expression data from both cohorts were combined into a single, unified count matrix with genes as rows and spatial segments (ROI × cell type) as columns. Segments matched with fully annotated metadata were retained. Raw gene counts were then normalized, and for the EBV-positive vs EBV-negative DLBCL dataset (**Fig. 4**), additional rigorous batch correction steps were adopted as described below.

#### Rationale for batch correction

Overall, GeoMx datasets often involve samples from multiple cohorts and experimental batches, each potentially introducing technical artifacts that can obscure true biological variation. In the context of our DLBCL patient cohort, where samples are derived from diverse sources, correcting for batch effects is critical to ensure that the observed differences in gene expression reflect underlying biology rather than technical or sample processing discrepancies. Batch correction methods help to remove these unwanted sources of variation while preserving genuine differences arising from biological conditions and cell types. This step is important for downstream analyses such as differentially expressed gene (DEG) analysis and gene signature validation, as it ensures that identified biomarkers and signatures are robust and not confounded by technical and other unwanted factors.

#### Normalization methods, negative control genes, and unwanted covariant factor preparation for batch correction

The standR (86) (v.1.9.3) pipeline was used for normalization and reducing patient-level batch effects using the RUV4 method. Two normalization methods were adopted, including log counts-per-million reads (CPM) via the logNormCounts function of scater package (v.1.28.0) and quantile normalization via geomxNorm function of standR. Batch effect correction was implemented via a grid searching strategy to optimize parameter combinations for minimizing individual patient-level variations (e.g. tissue sources) while retaining biological variations due to EBV condition and cell types. Five grids of the number of negative control genes (NCG) were selected: 1000, 2000, 3000, 4000, and 5000 via findNCGs function. The three grids of the number of unwanted factors (i.e. k-values) for the RUV4 method (87) were set to 1, 2, and 3 using the geomxBatchCorrection function. The result of each batch correction run was a normalized and adjusted expression matrix for DEG.

#### DEG parameter settings

Following batch correction, a two-step approach was employed to evaluate and refine DEG parameters. First, the suitability and effectiveness of batch correction strategies were assessed by examining their ability to produce biologically interpretable DEGs. To do this, pairwise comparisons were conducted between key cell populations of interest (e.g. tumor, CD4T, CD8T, and macrophage compared with endothelial cells, respectively) across different EBV status subsets (EBV-positive, EBV-negative, and combined). These contrasts aimed to reveal condition-dependent DEGs that are biologically meaningful.

Second, the DEG model parameters were optimized to recover cell-type-specific gene signatures robustly. DEG analyses were performed using a pipeline that integrated edgeR (48) (v.3.42.4) and limma (88) (v.3.56.2). The modeling framework allowed for the inclusion of weight matrices from RUV4 in the design matrix of the linear model as covariates. Four confounder sets were tested:

1. No confounders
2. One confounder if the k-value is equal to or greater than 1: one weight matrix from RUV4.
3. Two confounders if the k-value is equal to or greater than 2: two weight matrices from RUV4.
4. Three confounders if the k-value is equal to 3: three weight matrices from RUV4.

Additionally, each confounder set was tested with two scenarios: with and without controlling for cell-type abundance (i.e. including or excluding cell counts as a covariate). DEGs were then identified using moderated linear modeling (limma) and empirical Bayes shrinkage. Significance thresholds included an adjusted p-value threshold of 0.01. P-values were adjusted for multiple testing using the false discovery rate (FDR) method.

#### Benchmarking and Signature Validation

To systematically assess the combined influence of batch correction and DEG model parameters, all combinations (N = 540) of number NCGs, k-values for unwanted variation, EBV status subsets, confounder sets, and cell-type abundance adjustments were evaluated. The DEGs identified under each parameter setting were then evaluated against known cell-type-specific signatures. Signatures (**Supp Table 8**) included well-established lineage and function markers for CD4 T cells (89), CD8 T cells (89), macrophages (90, 91), and DLBCL tumor cells (92). Enrichment of known markers within each DEG list was assessed via hypergeometric tests, confirming whether the parameters chosen successfully recovered expected biological signatures.

#### VisiumHD data

The demultiplexed FASTQ output files from next-generation sequencing were used to map and quantify the human probes through the 10x Genomics Space Ranger v3.1.1 count pipeline. Manual alignment and tissue detection was first performed with 10x Genomics Loupe Browser v8.0.0 using the CytAssist image and the H&E stained microscope image. These images, together with the human transcriptome reference GRCh38, Visium probe set v2.0, and the FASTQ files, were input into the Space Ranger’s count pipeline. Due to varying ROI sizes in the tissue samples, unique molecular identifier (UMI) counts were normalized by the number of bins within each ROI, with a scaling factor of 10,000. Note that batch effect correction was similarly not performed for the analysis in **Fig. 1C**.

#### CosMx data

The acquired data was automatically uploaded onto the AtoMx spatial informatics platform, with the normalized transcript counts of each FOV generated in the platform, as well as image pre-processing and feature extraction, To identify single-cell features, a pre-trained neural network model Cellpose was used for segmentation (93). Single-cell RNA expression profiles were generated by counting transcripts of each gene falling within different segmented areas. Cells with fewer than 20 total transcripts were removed from downstream data analysis.

### SGCC Development Rationale

The spatial distribution of cell phenotypes in tissues provides vital insights into cellular interactions, functional states, and tissue microenvironment organization. Spatial autocorrelation, commonly quantified using metrics like Moran’s I or Geary’s C, is a well-established measure for evaluating the degree of similarity in values across spatially adjacent locations for a single signal (e.g. cell phenotype distribution pattern). However, these methods are limited in their ability to compute cross-correlation between two spatial signals, particularly in scenarios involving graph-based data structures. In addition, traditional correlation methods such as Pearson and Spearman correlation, while effective for linear or rank-based relationships, are inadequate for measuring spatial relationships between two graph signals. To address this gap, we introduce Spectral Graph Cross-Correlation (SGCC), a method that quantifies the similarity between two graph signals by analyzing and comparing their spectral components in the frequency domain.

SGCC addresses these limitations by leveraging the Graph Fourier Transform (GFT) to analyze graph signals in the frequency domain. The rationale for SGCC lies in its ability to:

#### 1. Extend beyond single-signal analysis

While spatial autocorrelation measures like Moran’s I evaluate the spatial coherence of a single signal, SGCC quantifies cross-correlation between two graph signals, capturing their spatial relationship in terms of complementarity or co-occurrence.

#### 2. Incorporate graph structure

SGCC operates directly on graph-structured data, integrating spatial adjacency information into the analysis. This allows it to adapt to both regular (e.g. pixel grids) and irregular (e.g. cell-cell adjacency) spatial graphs, ensuring an accurate representation of spatial relationships.

#### 3. Focus on k-bandlimited signals to study spatially organized structures

A k-bandlimited signal refers to a smooth and slow graph signal, which can be biologically defined as a spatially organized structure (19) (e.g. germinal center pattern in a reactive tonsil). Such signal can be effectively captured by first k Fourier modes (FM), which are eigenvectors of graph Laplacian to capture broad, large-scale patterns in the graph data, such as gradual and organized distributions. In contrast, high-frequency signals represent rapid, small-scale variations that often correspond to noise or localized fluctuations. By focusing on k-bandlimited signals, SGCC isolates biologically meaningful spatial relationships while minimizing the influence of noise. This approach ensures that the analysis highlights overarching spatial trends, such as how two cell types are distributed across tissue regions, rather than being confounded by random variations.

#### 4. Provide a quantitative and interpretable metric

SGCC calculates the cosine similarity of Fourier coefficients (FC) of first k FM, offering a robust and interpretable metric for spatial co-localization. This measure effectively captures the similarity of large-scale spatial patterns while accounting for the graph structure.

#### 5. Enable cross-sample comparisons

By standardizing spatial data into a pixel graph and ensuring all regions of interest (ROIs) are represented within the same linear space, SGCC allows for consistent and comparable measurements across multiple samples or conditions.

#### 6. Link spatial patterns to functional insights”

SGCC integrates spatial cross-correlation with functional analyses, enabling the identification of spatially dynamic genes associated with the spatial arrangement of specific paired cell phenotypes. By connecting spatial patterns to gene expression, SGCC provides a comprehensive view of how spatial organization influences cellular behavior and tissue function.

### SGCC Development

#### Binning cell phenotype data into a grid

Note that all the notations of matrices and vectors are bolded, and all the vectors are treated as column vectors in the following description. Given a set of spatial coordinates (*x*_*s*_, *y*_*s*_) for each cell *s*, the tissue area is discretized into a regular grid. Each bin (or cell of the grid) aggregates cells of various types. For each cell phenotype, a count is computed per bin, resulting in a cell phenotype-specific spatial map. This step converts a potentially irregular distribution of cells into a uniform representation suitable for graph construction. Specifically, a one-hot encoded matrix **C** is first constructed, where rows represent cells and columns correspond to cell phenotypes, with each element *c*_*s,r*_ set to 1 if the cell *s* belongs to cell phenotype *r*, and 0 otherwise, where *s* = 1, 2, …, *c*, and *r* = 1, 2, …, *m*. This matrix is then transformed into a bin-by-cell phenotype matrix **P**, where rows represent bins in the grid, columns correspond to cell phenotypes, and each element *p*_*i,r*_ indicates the count of cells of phenotype *r* within bin *i*, where *i* = 1, 2, …, *n*, and *n < c*. This transformation ensures that spatial cell phenotype distributions are uniformly represented across the grid for downstream graph-based analyses. Based on the benchmarking results in **Supp Figs. 3A & B**, the default grid size is set as 60 × 60.

#### k-nearest neighbor (KNN) graph construction

Given a binned grid containing *n* pixels, including their spatial coordinates and cell type phenotype counts, SpaGFT first calculates the Euclidean distances between each pair of pixels based on spatial coordinates. Subsequently, an undirected graph *G* = (*V, E*) is constructed, where *V* = {*v*_1_, *v*_2_, …, *v*_*n*_} is the node set corresponding to the *n* pixels, and *E* is the edge set. An edge *e*_*ij*_ exists between *v*_*i*_ and *v*_*j*_ in *E* if and only if *v*_*i*_ is the KNN of *v*_*j*_ or *v*_*j*_ is the KNN of *v*_*i*_ based on Euclidean distance, where *i, j* = 1, 2, …, *n*, and *i* ≠ *j*. Based on the benchmarking results in **Supp Figs. 3A & B**, the default *K* is defined as 400.

An adjacency binary matrix **A** = (*a*_*ij*_) is defined, where rows and columns represent the *n* pixels:

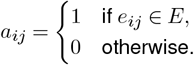

A diagonal degree matrix **D** = diag(*d*_1_, *d*_2_, …, *d*_*n*_) is then defined, where the degree of each node *v*_*i*_ is given by:

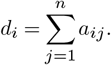

#### Fourier mode calculation

Using the adjacency matrix **A** and the degree matrix **D**, a Laplacian matrix **L** is defined as:

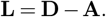

The Laplacian matrix **L** can be decomposed using spectral decomposition:

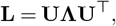

where **Λ** = diag(*λ*_1_, *λ*_2_, …, *λ*_*n*_) is a diagonal matrix containing the eigenvalues of **L**, ordered such that *λ*_1_ ≤ *λ*_2_ ≤ … ≤ *λ*_*n*_, and **U** = (***µ***_1_, ***µ***_2_,…, ***µ***_*n*_) is a matrix whose columns are the unit eigenvectors of **L**. Note that *λ*_1_ is always equal to 0, regardless of the graph topology, and is excluded from the subsequent analysis. Each eigenvector ***µ***_*k*_ corresponds to a Fourier mode (FM), where ***µ***_*k*_ ∈ ℝ^*n*^, *k* = 1, 2, …, *n*, and the set ***µ***_1_, ***µ***_2_,…, ***µ***_*n*_ forms an orthogonal basis for the linear space.

For 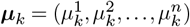, where 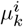 indicates the value of the *k*th FM on node *v*_*i*_, the smoothness of ***µ***_*k*_ reflects the total variation of the *k*th FM in all mutual adjacent nodes. This smoothness is formulated as:

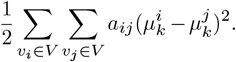

This expression can be derived using matrix operations:

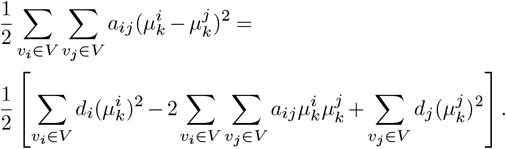

Simplifying further:

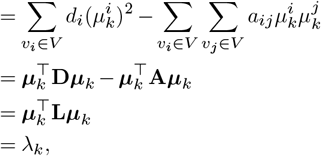

where 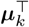 is the transpose of ***µ***_*k*_.

According to the definition of smoothness, a small eigen-value *λ*_*k*_ indicates a low variation in FM values between adjacent nodes, corresponding to low-frequency FMs. Conversely, larger eigenvalues correspond to higher oscillations in the eigenvectors, representing high-frequency FMs. Thus, the eigenvalues and eigenvectors of **L** are interpreted as frequencies and FMs in SpaGFT. Intuitively, low-frequency FMs capture broad, large-scale spatial patterns, while high-frequency FMs reflect finer, localized variations.

#### First k bandwidth determination by Kneedle algorithm

The eigenvalue *λ*_*t*_ is converted as follows:

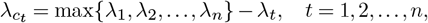

where 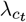 is the converted value of *λ*_*t*_. Each point 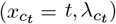, where 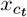 is the rank number of 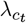, is processed by a smoothing spline to preserve the curve shape and obtain 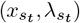, *t* = 1, 2, …, *m*. Denote the coordinate set as:

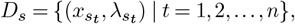

which can be normalized to the coordinate set *D*_*n*_ as follows:

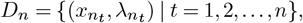

where:

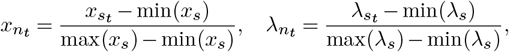

and min(*x*_*s*_), max(*x*_*s*_) are the minimum and maximum of 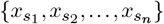, respectively. Similarly, min(*λ*_*s*_) and max(*λ*_*s*_) are the minimum and maximum of 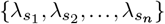, respectively. Additionally, let *D*_*d*_ represent the set of points corresponding to the differences between the *x*-and *λ*-values:

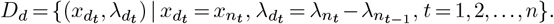

The determination of the cutoff *y*_*z*_ can then be converted to identifying the inflection point *λ*_*z*_, which satisfies:

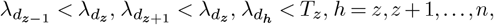

where:

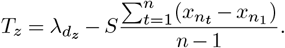

In the equation above, *S* is a coefficient that controls the level of aggression in identifying the inflection point; here, *S* is set to 2.

#### Graph Fourier Transform

The graph signal of a cell phenotype pattern *p* is defined as:

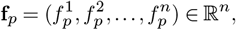

which is an *n*-dimensional vector representing the cell count values across *n* bins. The graph signal **f**_*p*_ is transformed into Fourier coefficients 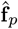 by:

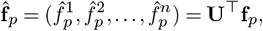

where 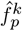 is the projection of **f**_*p*_ onto the *k*-th Fourier mode *µ*_*k*_, representing the contribution of *µ*_*k*_ to the graph signal **f**_*p*_, with *k* = 1, 2, …, *n*. This Fourier transform aligns the cell phenotype pattern with its spatial distribution, representing the pattern in the frequency domain.

#### SGCC calculation

After transforming the graph signals of two cell phenotype patterns **p**_·,1_ and **p**_·,2_ into their respective low-frequency representations, SGCC is computed by evaluating the cosine similarity of their *k*-bandlimited Fourier coefficients (FCs), capturing large-scale spatial distributions.

The SGCC score is calculated as:

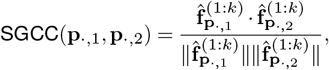

where:

- 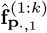 and 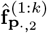 are the vectors of the first *k*-bandlimited FCs for cell phenotype patterns **p**_·,1_ and **p**_·,2_, respectively.
- 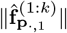 and 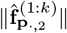 are the Euclidean norms of these coefficient vectors.

This measure yields a normalized similarity score between -1 and 1:

- A high SGCC score (close to 1) indicates that the two cell phenotypes exhibit similar large-scale spatial structures.
- A low or negative SGCC score (close to -1) suggests that the two cell phenotypes have inversely related spatial patterns at these scales.

For the IN-DEPTH data with *m* cell phenotypes, there are 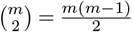 SGCC scores.

### SGCC Validation Analysis

#### Simulation 1 (ring pattern)

The simulation process begins by defining a regular 60 by 60 grid to represent the spatial domain, with each cell having x and y coordinates. An inner circle is generated with a fixed radius from a predefined range (2.5, 5, 7.5, 10, 12.5, 15, 17.5, and 20), centered in the middle of the grid (x=30, y=30). To simulate the dynamic behavior of an outer ring shrinking toward the inner circle, a sequence of radii is defined for the outer ring in 10 incremental steps, starting from a large initial radius and progressively decreasing to slightly larger than the inner circle’s radius. For each step, the grid is analyzed to classify points as either inside the inner circle, within the outer ring (defined as the area between the shrinking outer radius and the inner circle), or outside both regions. The spatial distribution of these classifications is aggregated for all steps, resulting in a set of data that captures the interaction between the inner circle and the shrinking outer ring at different stages of the simulation. This process enables the generation of 80 datasets to demonstrate local and global complementary patterns.

#### Simulation 2 (moving pattern)

The simulation method generates data to model the spatial interactions between two dynamically moving circular regions on a 60 by 60 grid. For each simulation, the radius of the first circle is varied within a specified range (6,7,8,9,10,11,12,13, and 14), while the radius of the second circle is set to be 1.5 times the radius of the first circle. Initially, the centers of the two circles are positioned symmetrically at a distance of 30 units from the centerline of the grid. Over 10 incremental steps, the centers of the circles move inward toward the grid’s center. At each movement step, the Euclidean distance from every grid point to the centers of the circles is calculated to determine whether a point lies within the first circle, the second circle, both circles or outside both. This classification is updated at each step to reflect the movement of the two circles. The resulting data for each simulation step includes the binary indicators for points being within each circle and the overlap between the two. This process enables the generation of 80 datasets to demonstrate moving pattern of two cell types.

#### Space-gene covarying analysis

To investigate spatially covarying gene expression in relation to cell-cell spatial pattern dynamics across multiple samples, SGCC scores are leveraged as spatial factors and treated as time-series variables within the ImpulseDE2 framework (v0.99.10). ImpulseDE2 is a statistical tool designed for differential expression analysis, employing a sigmoid-based impulse model to represent continuous trends across time. By utilizing SGCC scores as a continuous spatial variable, this approach facilitates the identification of genes whose expression systematically correlates with spatially defined paired cell phenotype patterns, enabling the exploration of underlying molecular mechanisms associated with changed spatial organization across multiple samples or ROIs.

The workflow begins by addressing batch effects using previously established batch correction methods (as detailed above and also in (21)). Following this, the input consists of a gene expression matrix, sample metadata, and SGCC scores, which represent the spatial relationships between paired cell phenotypes. The dataset is preprocessed by subsetting to include relevant cell phenotypes and experimental conditions while correcting for batch factors using default ImpulseDE2 settings. In **Fig. 3E**, CD4T cells and BCL6-positive B cells were selected. If metadata is available, it is constructed for each sample, incorporating binary conditions (e.g. case vs. control), SGCC scores as continuous spatial factors, and batch information. SGCC scores are then discretized into time bins to represent progression along the spatial factor for time-series modeling. Using ImpulseDE2, a sigmoid-based impulse model is applied to capture non-linear gene expression dynamics across SGCC-defined time bins. Genes are ranked based on their temporal expression trends and categorized into patterns such as increasing, decreasing, or transient, and significant genes are identified using an adjusted p-value threshold based on the Benjamini–Hochberg (BH) method. The output consists of a ranked list of genes that covary with the spatial factor, classified patterns of gene expression, and insights into spatially regulated molecular mechanisms linked to changes in paired cell phenotypical patterns.

### Spatial Proteomics Analysis

#### Image processing

For functional markers included in the analysis in **Fig. 4** (HLA-1, HLA-DR, CD45RO, CD45RA, Ki-67, PD-1, LAG3, Granzyme B), the 16-bit intermediate QPTIFFs, generated by the Phenocycler Fusion, were used to ensure optimal dynamic range of data. The QP-TIFFs were processed firstly by subtracting the last blank cycle scaled by the ratio between current channel cycle and total cycle number, i.e.,

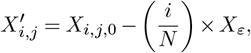

where 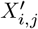 is the blank-subtracted image of marker *j* in cycle *i*; *X*_*i,j*,0_ is 16-bit intermediate image of marker *j* in cycle *i*; and *X*_*ε*_ is the last blank cycle. Then, the last-blank-subtracted image were processed in imageJ using the “Math” and “Subtract Background” functionalities under “Process”:

1. Subtract the mean pixel value of the image to get rid of most of the “salt and pepper” noise.
2. Subtract the background generated by the sliding paraboloid algorithm with a 5 pixel radius.

Since GeoMx images were outputted as 16-bit images by default and were already fully processed internally by the instrument, Tox and PD-L1 were not processed by the above-mentioned pipeline. Finally, for each core and each marker, a lower bound and an optional upper bound (in case of high pixel intensity artifacts) were applied to remove the remaining unspecific staining, noise, and artifacts. The lower bound and upper bound were determined by visual inspection of the images in QuPath and the values can be found in **Supp Table 9**.

Note that cell phenotyping was performed based on the final 8-bit QPTIFF generated by the Phenocycler Fusion. Since the 8-bit QPTIFF was processed completely by the Phenocycler Fusion’s software, the blank subtraction and the imageJ processing were not applied. However, similar to the 16-bit images, lower bounds were set for each core and each marker in order to get rid of as much of unspecific staining (for example, nuclear signal of a supposedly membrane marker) as possible. The lower bound values can be found in **Supp Table 9**.

#### Data processing

The aforementioned functional markers (HLA-1, HLA-DR, CD45RO, CD45RA, Ki-67, PD-1, LAG3, Granzyme B, Tox, PD-L1), were scaled by the respective median nuclear signal (DAPI for markers captured on Fusion and SYTO13 for markers captured on GeoMx) of each tissue sample in order to adjust for different binding efficiency of markers. Then, a global min-max scaling was applied to scale the marker expression levels to be within [0,1].

For phenotyping markers (Pax5, CD20, CD3, CD8, CD4, FoxP3, CD11c, CD68, CD163, CD31), the same median nuclear signal scaling was applied. Then, the markers were further scaled within each tissue sample by a (0.001, 0.999) quantile scaling and then truncated at 0 and 1. Unlike the functional markers, the phenotyping markers were scaled at a local level to compensate for tissue samples with an overall weaker pixel intensity.

#### Marker enrichment heatmap

The marker enrichment heatmap showed the Z-score of a given (marker, cell type, EBV status) tuple. In other words, it showed how many standard deviations away is the mean of marker A expression of cell type B given an EBV condition from the population mean of marker A expression:

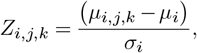

where *Z*_*i,j,k*_ stands for the Z-score for marker *i*, cell type *j*, and EBV status *k*; *µ*_*i,j,k*_ stands for the mean expression for for marker *i*, cell type *j*, and EBV status *k*; *µ*_*i*_ stands for the population mean of marker *i*; and *σ*_*i*_ stands for the population standard deviation of marker *i*.

#### Cell type proportion and enrichment

Cell type enrichment was presented as log_2_ of the ratio between the proportion of cell types in EBV-positive and EBV-negative DLBCL samples:

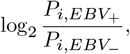

where 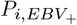 is the proportion of cell type *i* in EBV-positive and 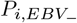 is the proportion of cell type *i* in EBV-negative.

#### Dysfunction score

The T cell dysfunction score constructed to measure the overall dysfunction of a cell includes markers that are differentially expressed. PD-1 was not included due to its lower staining quality in this tissue cohort, as well as its additional biological function as an activation marker (94).

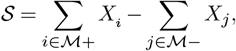

where 𝒮 stands for the dysfunction score; *X*_*i*_ and *X*_*j*_ stands for the expression level of marker *i* or marker *j* of a cell; ℳ+ stands for a set of markers that signify contributive effects to cell dysfunction, ℳ+ = {LAG3, CD45RO, Tox}; ℳ− stands for a set of markers that signify counteractive effects to cell dysfunction, ℳ− = {CD45RA, Ki67, GZMB}.

#### Cell motif analysis

For a tissue sample, each cell’s spatial location was recorded as the (x,y) of the centroid of its segmentation mask. Using the set of centroids, a Delauney triangulation was first performed. Then a graph was constructed using the simplices. Two nodes were connected if and only if the Euclidean distance between the two nodes is less than or equal to 20um. For each node of interest, for example, all CD4 T cell nodes, its immediately adjacent nodes, i.e. one-hop neighbors, were identified. Then, the composition of a given one-hop neighborhood was summarized into a vector representing the count of each cell type. For example, a one-hop neighborhood might consist of 2 CD4 T cells and 1 CD8 T cells, while there were 4 annotated cell types in total, the summary vector would be (2, 1, 0, 0). These vectors were then clustered using K-means clustering to find repeating motifs.

#### Negative binomial regression

Two negative binomial regression models were fitted to explore the effect of EBV status, membership of motif, and their interaction on M1-like macrophage and M2-like macrophage counts within the one-hop neighborhood anchoring on CD4 T cells. The proposed model is:

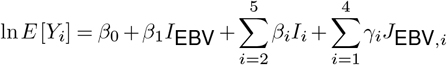

where

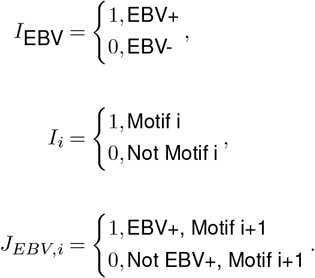

#### Tumor density score

Tumors were first classified into three categories:

- EBV-positive, LMP1 high: if a tumor is in an EBV-positive sample and its LMP1 expression is greater than the median LMP1 expression of all tumors.
- EBV-positive, LMP1 low: if a tumor is in an EBV-positive sample and its LMP1 expression is less than or equal to the median LMP1 expression of all tumors.
- EBV-negative: if a tumor is in an EBV-negative sample.

Tumor density score was then calculated as described in (21). Briefly, within each of these categories, for each non-tumor cell, three tumor scores were calculated, one for each tumor class. The score was calculated based on a cell’s distance to tumors within a closed neighborhood of radius *r*. Let **J** = {1, …, *m*}denote the indices of all the tumors in the dataset and *d*_*i,j*_ denote the distance from the cell *i* to tumor *j*. Then, the tumor score is calculated as

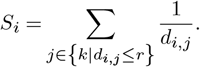

Then, the score was transformed into

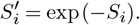

### Spatial Transcriptomics Analysis

#### RNA quantity comparison

The non batch-corrected CPM counts (GeoMx data), UMI counts (VisiumHD data), and transcript counts (CosMx data) were used as gene expression measurements after log1p transformation. Pearson correlation coefficients were calculated for each adjacent IN-DEPTH and control slide pairs, with each datapoint being 1 unique gene. Total RNA quantity, as well as total control RNA quantity, were generated by first summing all the respective gene counts across the ROIs, and then visualized on a log1p scale. Genes labeled as “NegProbe” or “Neg” in the GeoMx and CosMx probe kits were used to determine the control probe counts; note that the VisiumHD probe panel did not include any internal negative controls.

#### Gene signature curation and scoring

All gene signatures used in this study (95), apart from those that were manually curated, were obtained using the R package ‘msigdbr’ (v7.5.1), and the enrichment of gene signatures within cell populations were calculated using Gene Set Variation Analysis (GSVA) (96) through the R package “gsva” (v1.52.3) with the default parameters.

The gene signatures used to validate the transcriptomic signature of annotated cell populations (**Fig. 2C, middle**) were were derived from a tonsil scRNAseq atlas comprising over 556,000 cells (35). They were used to (1) calculate cell type associated differential expressed genes (DEG) for enrichment analysis of IN-DEPTH captured transcriptomics data, and (2) provide scRNA-seq reference for deconvolution analyses. The processing workflow began by loading Seurat objects (97) (v4.4.0). Cells were subsampled and refined to merge to reduce dataset complexity based on the annotation with 135 cell types. Specifically, “SELENOP FUCA1 PTGDS macrophages,” “C1Q HLA macrophages,” “ITGAX ZEB2 macrophages,” and “IL7R MMP12 macrophages” were assigned as M2-like macrophages, “Mono/Macro” and “cycling myeloid” were assigned as myeloid cells. Cell types unrelated to this study, such as “cycling FDC,” “cycling T,” “granulocytes,” “DN,” “Granulocytes,” “ILC,” “Mast,” “NK,” and “preB/T,” were excluded from the analysis. The major B cell populations, including naive B cells (NBC), memory B cells (MBC), and germinal center B cells (GCBC), were refined by removing corresponding cell subsets with fewer than 100 cells. Overall, NBC, MBC, GCBC, CD4 T cell, CD8 T cell, Treg, M2-like macrophages, M1-like macrophages, myeloid, dendritic cell (DC), and epithelial cells were refined and extracted for enrichment and deconvolution analyses. Note that endothelial signatures were collected separately (98). Additionally, the Tfh signature used in **Fig. 2E** was curated using all unique genes from four annotated Tfh populations (“Tfh TB border”, “Tfh-LZ-GC”, “GC-Tfh-SAP”, “GC-Tfh-OX40”) in the same atlas resource (35).

DEG analysis was subsequently performed using Seurat (97) (v4.4.0) to identify gene signatures associated with specific cell types. Followed by the log-count-per-million (LogCPM) normalization method, the “FindMarkers” function was applied with default parameters, including a log fold-change threshold (log2FC > 0.25) and an adjusted p-value threshold (p adj < 0.05). For each cell type, DEGs were calculated by comparing the target cell population to all other cell types. Specifically, DEGs of NBC, MBC, GCBC, CD8 T cells, DC, and epithelial cells were identified by comparing each cell type with other cell types. DEGs of CD4 T cell and Treg by comparing each other. DEGs of M2 macrophage was compared with M1 macrophage. GSVA (96) (v.1.52.23) was used to determine enrichment of each gene signature (**Fig. 2C**). All gene signatures used in **Figs. 2C & 2D**, for tonsil cell types and Tfh cells, are in **Supp Table 1**.

The source and full names for gene signatures across **Figs. 3, 5** and **Supp Fig. 7** are in **Supp Fig. 3E**. The RNA gene signature for T cell dysfunction (**Fig. 4G, right** and **Fig. 4J, right**) was curated using a panel of genes that were previously described to be markers expressed on dysfunctional exhausted CD4 and CD8 T cells (51, 52, 99–101): *CTLA4, HAVCR2, LAG3, PDCD1, BTLA, TIGIT, CD160, CD244, ENTPD1, VSIR*. The EBV score RNA gene signature in **Fig. 5A** was generated using the average normalized counts for each detected EBV gene: *EBER1, EBER2, EBNA1, EBNA2, EBNALP, LMP1, RPMS1, BALF1, BCRF1, BHRF1, BNLF2A, BNLF2B, BNRF1, BZLF1*, with the expression of each EBV transcript also shown in **Supp Fig. 7A, top**.

#### Lymphocyte spatial distribution

The follicle-high and follicle-low regions were visually identified, with ROIs 3, 5, 17 from both tissues used for the former, and ROIs 1, 7, 14 from both tissues used for the latter (**Supp Fig. 2B**) to generate 6 data points for each follicle regions, after which the CD4 T cell Tfh GSVA scores were compared between these two follicle regions. Tfh correlation was determined by performing a Spearman correlation across all ROIs between each ROI’s B-cell proportion and CD4 T cell Tfh GSVA score.

#### Gene expression program (GEP) identification

GEPs were identified using consensus non-negative matrix factorization (cNMF) (44). The number of highly variable genes to use for cNMF was determined by setting a minimum threshold of 10% of all genes (at least 1800 genes in this case). The variance for all genes was then determined using the “FindVariableFeatures” function in Seurat (v4.4.0) (97), followed by k-means clustering with 9 centers with the random seed 1, to identify the cluster with the optimal cutoff for the number of highly variable genes. The number of genes chosen was then rounded up to the nearest hundred and used for cNMF. A range of 25 to 30 components (also known as GEPs) was tested for cNMF, an empirically determined optimum based on prior experience. The number of components with highest stability, where the stability is larger than the error, was chosen; in this case it was 26. The R package ‘enrichR’ (v3.2) (102) was then used to infer the biological function of each GEP by referencing the top 5 enriched GO Biological Process (GOBP) gene signatures (**Supp Table 2**). GEPs with at least 1 statistically significant (padj < 0.05) GOBP signature were determined to be distinctly enriched and were annotated based on their significant GOBP terms. The annotatable GEPs were then used to determine their relative enrichments across all the tonsil cell subpopulations in **Fig. 2F**.

#### Macrophage M1/M2 polarization and T cell dysfunction

Within each ROI, the proportion of M1-like and M2-like macrophages was calculated by (M2/(M1+M2)). To determine M2-rich and M1-rich subpopulations, the distribution of M2-like macrophage proportion was first plotted. The intersection of EBV-positive and EBV-negative distributions was then identified using the R package ‘pracma’ (v2.5.5), and was used to assign ROIs into the respective M1-rich and M2-rich subpopulations. Analysis on T cell dysfunction was then performed on the corresponding CD4 and CD8 T cell populations using the T cell RNA dysfunction signatur as described above.

#### CosMx cell phenotyping and analysis

Seurat (v4.4.0) (97) was used to perform unsupervised clustering and annotation of single cells. Harmony (v1.2.0) (103) was used for batch effect correction across different FOVs. Afterwards, the read count for each gene was divided by the total gene counts within each cell, multiplied by a scale factor of 100,000, and natural-log transformed. Principal component analysis (PCA) was performed on the normalized expression matrix using 2,000 highly variable genes. The top 15 principal components (PCs) were selected with a resolution parameter equal to 1. The clustering results were visualized using Uniform Manifold Approximation and Projection (UMAP) (104). We annotated cells into 5 major types according to their marker genes: *CD3D, CD4, CD8A* for T cells, *CD79A, MS4A1, MZB1, JCHAIN* for B/Plasma cells which were re-annotated as tumor cells, *LYZ, CD68, C1Q* for myeloid cells, *COL1A1, ACTA2* for fibroblasts, and *VWF, PECAM1, ENG* for endothelial cells. Note that batch correction was only performed for the analysis in **Fig. 5E**. Afterwards, GSVA (96) (v.1.52.23) was used to calculate T cell dysfunction signature enrichment in the annotated T cell population.

### Benchmarking of Deconvolution Softwares

#### CIBERSORT

CIBERSORT (40) is a computational method designed for cell type deconvolution from bulk tissue gene expression data using a reference-based approach. It employs a support vector regression framework (nu-SVR) to estimate cell proportions within a mixed tissue sample. The input includes a gene expression reference matrix, derived from the create_profile_matrix function of SpatialDecon, and a bulk tissue expression matrix in raw count format, created by combining and merging data across regions of interest (ROIs). The method is executed using the cibersort function, with parameters specifying the reference matrix and bulk expression data, enabling a robust deconvolution process that accurately quantifies cell type proportions.

#### dtangle

dtangle (41) (v2.0.9) is another method based on single-cell reference data that uses a linear scoring approach to estimate cell type proportions in bulk tissue samples. The input consists of a bulk tissue expression matrix and a single-cell dataset, both preprocessed to retain the most informative genes and cell types. The function dtangle facilitates the deconvolution by specifying parameters such as the combined dataset, the number of markers to use, and the data type. This ensures precise estimation of cell type proportions while maintaining compatibility with bulk and single-cell data formats.

#### MuSiC

MuSiC (42) leverages single-cell reference data for cell type deconvolution in bulk gene expression profiles. It employs weighted non-negative least squares to estimate the contributions of distinct cell types within bulk samples. The input includes the same bulk expression matrix used in CIBERSORT and a single-cell expression dataset formatted as a SingleCellExperiment object. This dataset is preprocessed to include cell types of interest and differentially expressed genes to enhance deconvolution accuracy. The deconvolution process is implemented through the music_prop function, where users specify key parameters, including cell type annotations and sample identifiers, ensuring the alignment of single-cell and bulk datasets.

#### SpatialDecon

SpatialDecon (43) (v1.13.2) utilizes a lognormal regression model to perform gene expression deconvolution. Unlike other tools, it can integrate normalized bulk expression data and single-cell reference matrices. The method aligns genes across datasets to ensure consistency during deconvolution. The spatialdecon function allows users to specify the normalized bulk expression data, background adjustment parameters, and the reference matrix. This method is particularly effective in leveraging both single-cell and bulk datasets to provide accurate cell type proportion estimates, while the alignment step enhances consistency across data sources.

### Application of SGCC on DLBCL Dataset

To analyze DLBCL GeoMX data, we first calculated SGCC scores to capture spatial relationships between the cell phenotypes. Samples were merged and discretized into a uniform 60 by 60 bin grid. Pairwise SGCC scores were computed for all cell types, reflecting their large-scale spatial distributions.

For DEG analysis between EBV-positive and EBV-negative conditions, we applied edgeR (48) and limma frameworks with batch corrected data (batch correction performed as described in the Batch Correction section). Batch corrected data were fitted to a linear model using the “mFit” function, incorporating a pre-defined design matrix. Empirical Bayes moderation was applied using the “eBayes” function to stabilize variance estimates, followed by DEG identification with the “topTable” function, ranked by adjusted p-values. Specific normalization strategies and batch correction parameters were applied based on cell types:

- CD4 T cells: LogCPM normalization, top 5000 NCGs, k=2, using two weight matrices from RUV4 batch correction, with cell type number included as a covariate in the design model.
- Macrophages: LogCPM normalization, top 1000 NCGs, k=3, using three weight matrices from RUV4 batch correction as covariates.
- Tumor cells: LogCPM normalization, top 1000 NCGs, k=3, using one weight matrix from RUV4 batch correction as a covariate.

DEGs between EBV-positive and EBV-negative conditions for CD4 T cells, macrophages, and tumor cells were filtered based on adjusted p-value thresholds (padj < 0.01, BH method). Enrichment analysis was performed for each DEG set using the enrichR (102) (v3.2) database, focusing on “Reactome_2022,” “GO_Biological_Process_2023,” and “KEGG_2021_Human”. Genes enriched in biologically-meaningful pathways (**Fig. 5, Supp Fig. 7**, and **Supp Table 2**) were selected for GSVA analysis to refine functional insights. Heatmap visualization was subsequently generated to highlight pathway activity across conditions based on ComplexHeatmap (v2.16.0). ggtern (v3.5.0) was used for visualizing CD4 T cell, Tumor, and Macrophage ternary plots using SGCC scores from CD4 T cell-Tumor, Macrophage-Tumor, and CD4 T cell-Macrophage (**Supp Table 4**). The adjacency enrichment statistic (AES) for each cell pair was determined as described in (54), where the expected number of edges between cell types was computed based on the frequencies of the cell types and the total number of edges in the graph. Specifically, AES was then calculated by comparing the observed number of edges connecting the two cell types to the expected number of edges. An AES of 0 indicates no enrichment over expectation, while positive and negative values indicate enrichment and depletion, respectively. Additionally, the density transparency was mapped to contour levels and color-coded by EBV status (i.e. “EBV+” and “EBV-”).

## DATA AVAILABILITY

## CODE AVAILABILITY

## ACKNOWLEDGEMENTS

We thank Craig Lassy and Michael Hair from Akoya for Phenocycler Fusion technical support, and Marvin Nayan, Adam Limb, Mike Chen, Brendan Collins, Nicholas Merino, Clement David, Sarah Miseirvitch, Prajan Divakar, Ozge Getkin, Tim Riordan, and Sarah Weigel from Nanostring for technical support. We also thank Jixin Liu, Jim DeCaprio, and other members of the Jiang and Ma labs for insightful discussions.

S.J. is supported in part by NIH DP2AI171139, P01AI177687, R01AI149672, R01GM152585, U24CA224331, a Gilead’s Research Scholars Program in Hematologic Malignancies, a Sanofi iAward, the Dye Family Foundation, the Broad Next Generation Award, and the Bridge Project, a partnership between the Koch Institute for Integrative Cancer Research at MIT and the Dana-Farber/Harvard Cancer Center. Q.M. is supported in part by NIH R01GM152585, P01CA278732, P01AI177687, U54AG075931, R01DK138504, and the Pelotonia Institute of Immuno-Oncology (PIIO). S.J.R. is supported by a Blood Cancer Discoveries Grant Program from the Leukemia Lymphoma Society, and The Paul G. Allen Frontiers Group. Y.Y.Y. is a recipient of the Albert J Ryan Fellowship. S.P.T.Y is a MacMillan Family Foundation Awardee of the Life Sciences Research Foundation.

This article reflects the views of the authors and should not be construed as representing the views or policies of the institutions that provided funding.

## AUTHOR CONTRIBUTIONS

Conceptualization: S.J., Y.Y.Y., Y.C., Q.M.

Experiment: Y.Y.Y., S.P.T.Y., H.A.M., H.Q.

Analysis: Y.C., H.Q., Y.Y.Y.

Contribution of reagents, tools, or technical expertise: W.W., X.J., S.K., L.P., S.Luo., P.C., J.L.L., Y.W., J.Y., N.E.A., B.S., R.M., M.V.O., W.L., K.J.L., S.Li., J.S., L.K., R.N.A.H., S.N., D.N., S.Sad., P.R., L.F., L.K., B.Zhu., A.B., N.D., C.N.C., J.K., Y.W.C., C.Y.C., J.Y.J.L., H.W., B.Zhao., L.H.L., D.M.K., V.B., B.Zhang., A.K.S., B.H., S.Sig., C.M.S., F.S.H., W.R.B., S.J.R.

Writing: Y.Y.Y., Y.C., H.Q., S.J., with input from all authors. Supervision and funding: S.J., Q.M.

Y.Y.Y., Y.C., and H.Q. contributed equally and have the right to list their names first in their C.Vs.

