## Supplementary Data for "Same-Slide Spatial Multi-Omics Integration Reveals Tumor Virus-Linked Spatial Reorganization of the Tumor Microenvironment"

[1]Center for Virology and Vaccine Research, Beth Israel Deaconess Medical Center, Harvard Medical School, Boston, MA, United States
[2]Program in Virology, Division of Medical Sciences, Harvard Medical School, Boston, MA, United States
[3]Pelotonia Institute for Immuno-Oncology, The Ohio State University, Columbus, OH, United States
[4]Department of Biomedical Informatics, College of Medicine, Ohio State University, Columbus, OH, United States
[5]Department of Medical Oncology, Dana-Farber Cancer Institute, Harvard Medical School, Boston, MA, United States
[6]Department of Pathology, Brigham and Women's Hospital, Harvard Medical School, Boston, MA, United States
[7]Translational Immunogenomics Lab, Dana-Farber Cancer Institute, Harvard Medical School, Boston, MA, United States
[8]Department of Pathology, Stanford University School of Medicine, Stanford, CA, USA
[9]Department of Pathology and Laboratory Medicine, University of Rochester Medical Center, Rochester, NY, United States
[10]Department of Pathology and Neuropathology, University Hospital and Comprehensive Cancer Center Tübingen, Tübingen, Germany
[11]Broad Institute of Harvard and MIT, Cambridge, MA, United States
[12]Department of Oral Medicine, Infection and Immunity, Harvard School of Dental Medicine, Boston, MA, United States
[13]Bioinformatics Institute (BII), Agency for Science, Technology and Research (A*STAR), Singapore, Singapore
[14]Department of Medicine, Brigham and Women's Hospital and Harvard Medical School, Boston, MA, United States
[15]Department of Hematology Oncology, Beth Israel Deaconess Medical Center, Harvard Medical School, Boston, MA, United States
[16]Department of Pathology, Dana Farber Cancer Institute, Boston, MA, United States
[†]Co-first authors
[#]Co-second authors
[*]Senior authors

#### Supplementary Figures

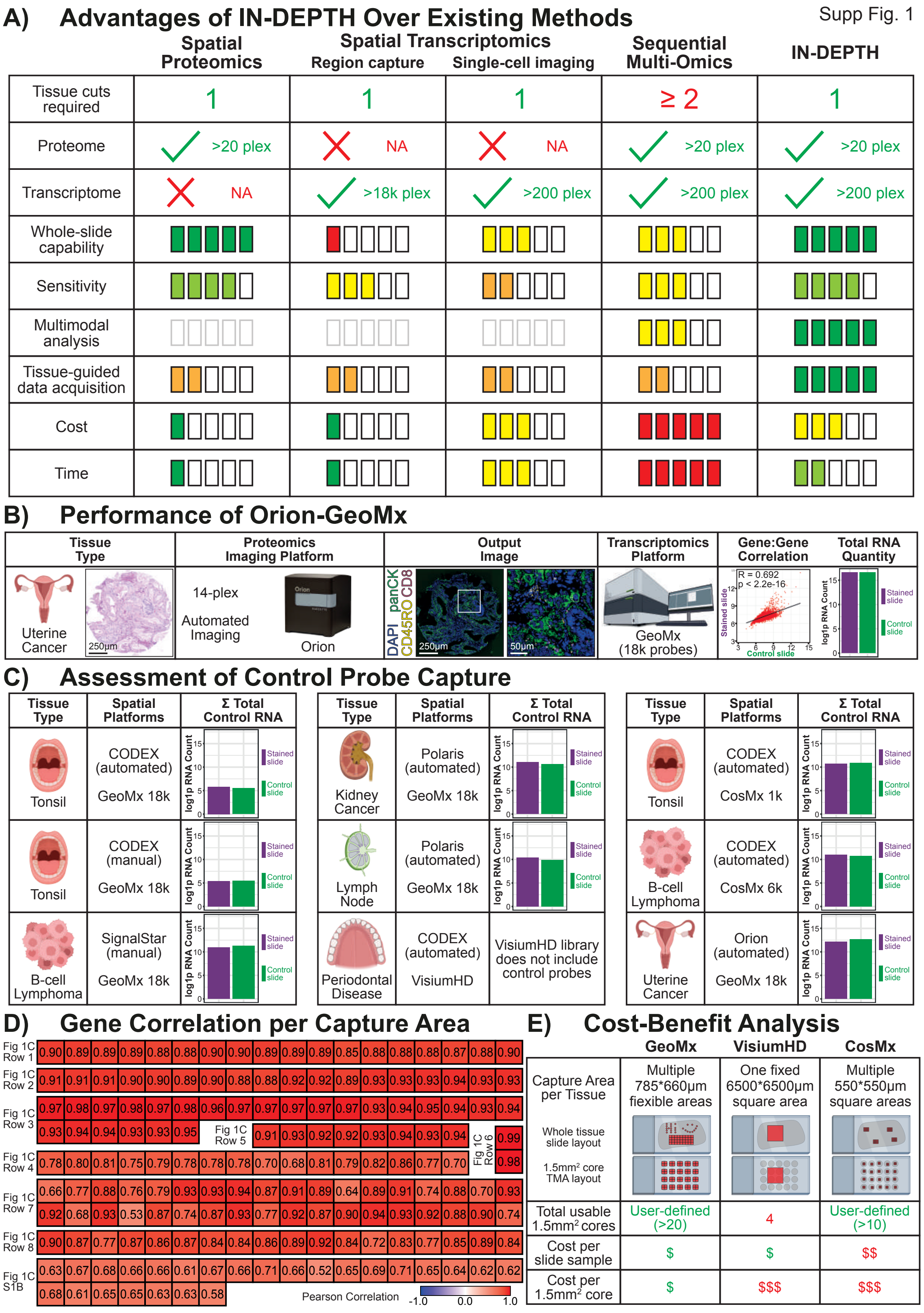


**Supplementary Figure 1: Breakdown of the advantages and additional quality control analysis for IN-DEPTH. (A)** Table comparing IN-DEPTH and other existing spatial-omics approaches. **(B)** Assessment of tissue imaging and RNA capture quality after Orion-GeoMx on uterine cancer tissues. **(C)** Comparison of total non-binding control RNAs captured across each IN-DEPTH experiment. **(D)** Breakdown of gene-to-gene Pearson correlation for each individual profiled ROI. **(E)** Table summarizing a cost-benefit analysis for IN-DEPTH with GeoMx, Visium HD, or CosMx as a transcriptome readout.

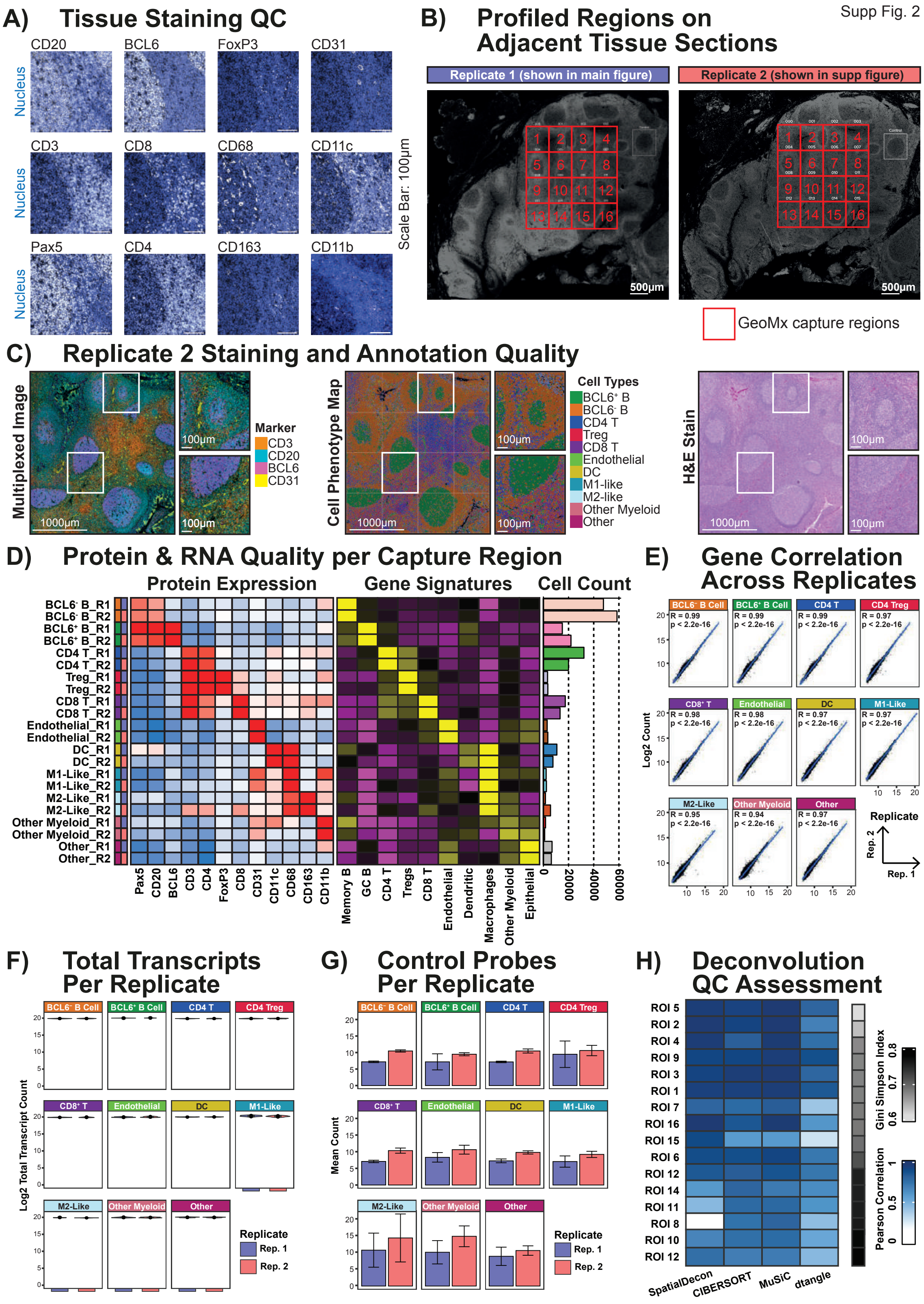


**Supplementary Figure 2: IN-DEPTH reproducibility across serial tissue sections using independent GeoMx instruments. (A)** Individual marker validation showing representative CODEX staining quality and distribution for each antibody in the 12-marker panel. **(B)** Full-slide image showing the adjacent tonsil tissues profiled, with the 16 spatially registered capture regions (highlighted in red) used for reproducibility assessment. **(C)** Technical reproducibility showing multiplexed protein staining, cell type annotation, and H&E imaging for the replicate tissue section presented in **Fig. 2B**. **(D)** Comprehensive comparison between replicates demonstrating consistency in protein expression patterns (left), cell type-specific transcriptomic gene signatures (middle), and cell counts (right) for each tissue replicate. **(E)** Gene-to-gene Pearson correlation analysis between replicates for each identified cell population to validate transcriptional reproducibility. **(F)** Total count of captured transcripts for each cell population across each tissue replicate. **(G)** Technical quality control showing consistent background signal levels through negative control probe counts across replicates. **(H)** Assessment of deconvolution accuracy by calculating the Pearson correlation between the computed cell proportions from each deconvolution algorithm and the IN-DEPTH-derived ground truth measurements across 11 cell types for each ROI. Cell type proportion complexity of each ROI is calculated using Gini-Simpson Index. All ROIs were ranked by the Gini-Simpson Index from low to high (top to bottom), indicating cell type proportion complexity from low to high.

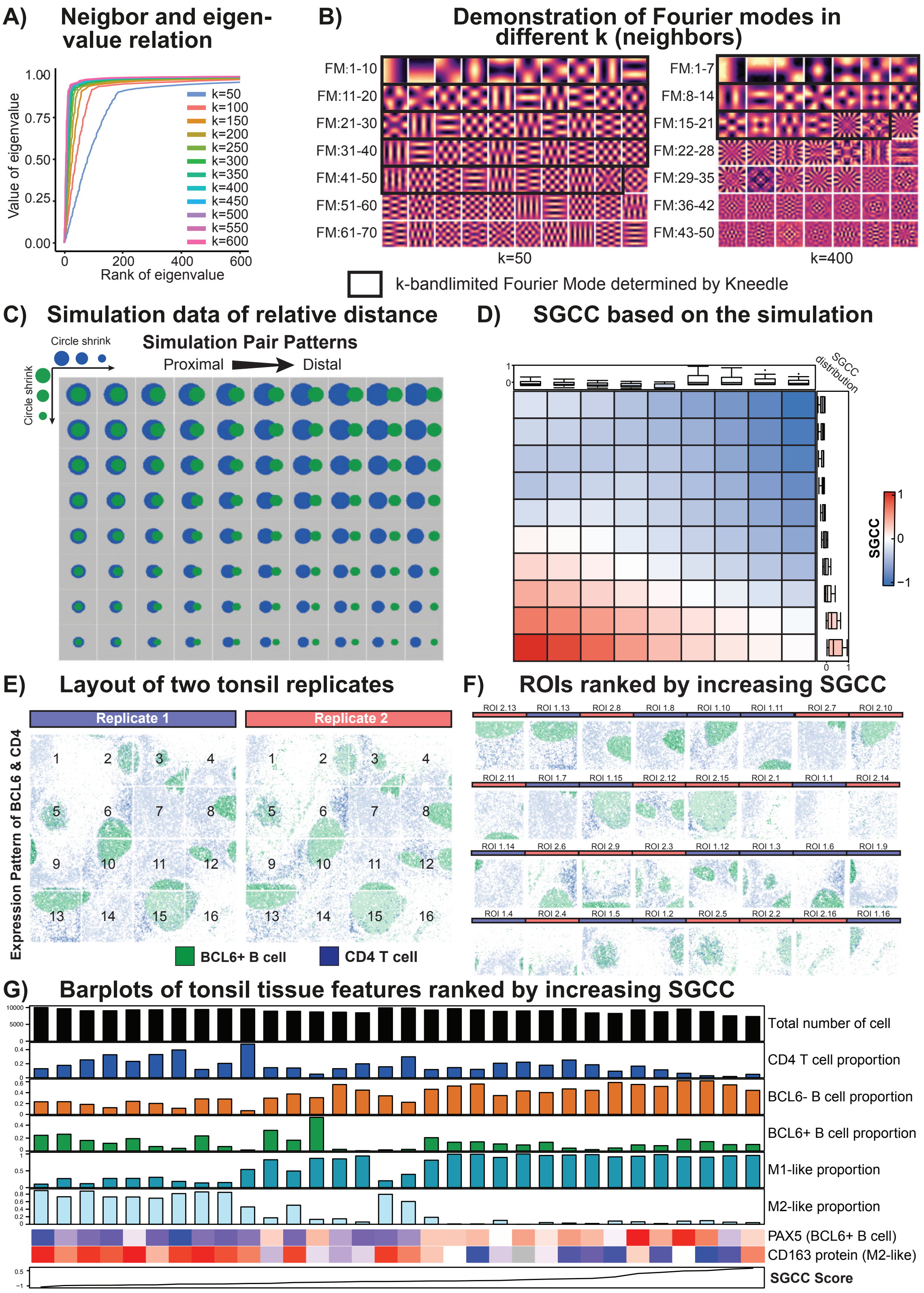


**Supplementary Figure 3: Validation and parameter optimization for SGCC analysis. (A)** The figure demonstrates a graph smoothness test based on different numbers of neighbors in constructing a kNN graph with 3600 nodes, where the x-axis is an eigenvalue of the laplacian matrix derived from the kNN graph; the y-axis is the value of the eigenvalue, showing the graph smoothness (the lower value corresponds to more smoothness). Lines are colored by the number of neighbors when constructing the kNN graph. **(B)** Visualization of patterns captured by k-bandlimited FMs at k = 50 and k = 400. **(C)** Extended validation of simulated SGCC patterns using proximal to distal patterns of two cell phenotypes. **(D)** SGCC scores related to the proximal and distal pattern simulations in (C). **(E)** Visualization of CD4 T cells and BCL6-positive B cells across replication tissue sections with ROI annotations. **(F)** Ranking of SGCC scores of each ROI from each replicate from low to high, and their associated ROI visualization of CD4 T cells and BCL6-positive B cells. **(G)** Quantification of cellular composition, cell populations, and protein expressions associated with the increase in SGCC score, corresponding to the transitions shown in **Fig. 3E**.

### Tissue Staining QC for Each TMA Cohort



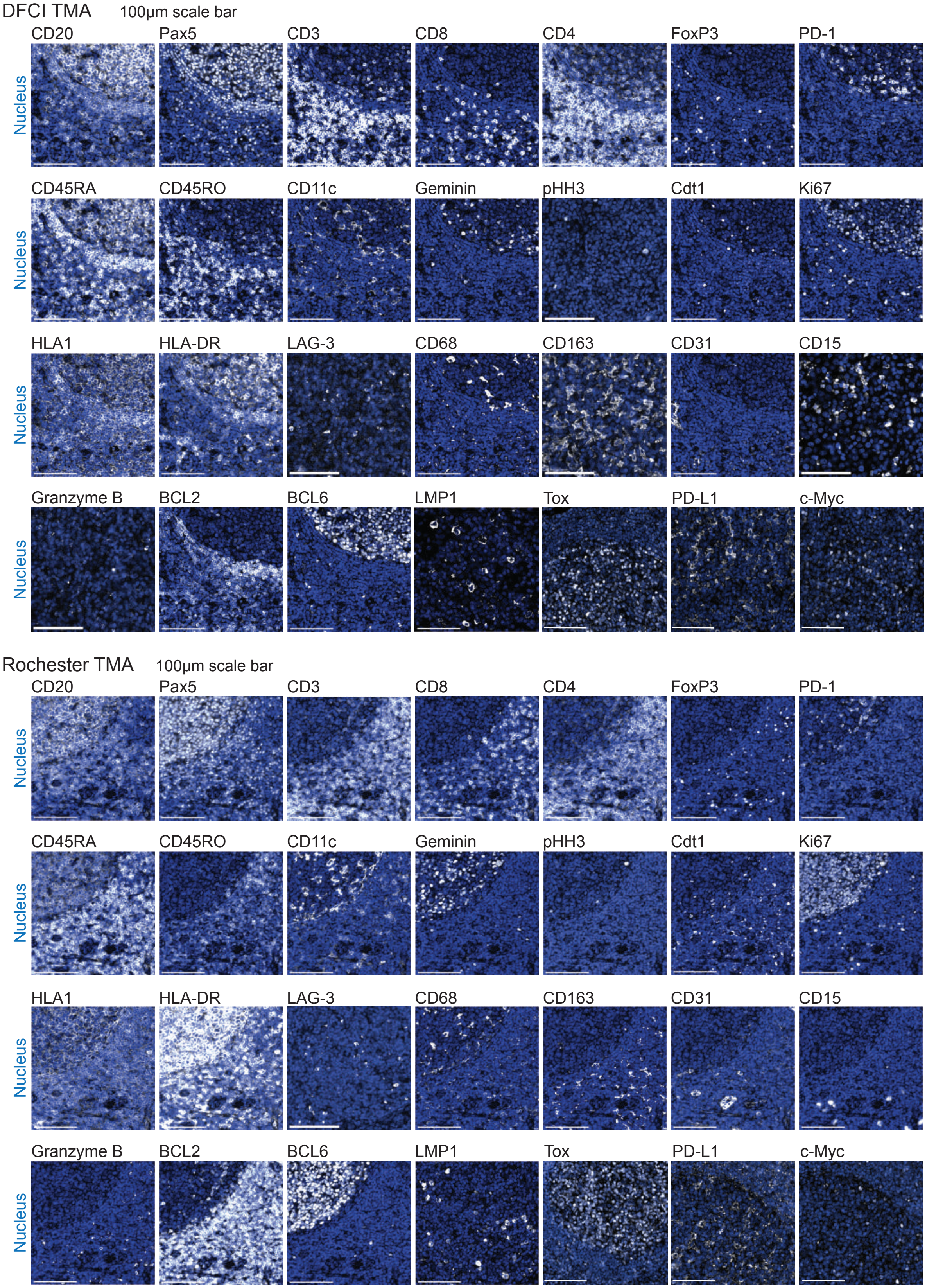


**Supplementary Figure 4: Validation of CODEX antibody staining specificity.** Representative CODEX images across DLBCL tissue sections, with the antibody markers (white) overlaid with cell nucleus stain (blue).

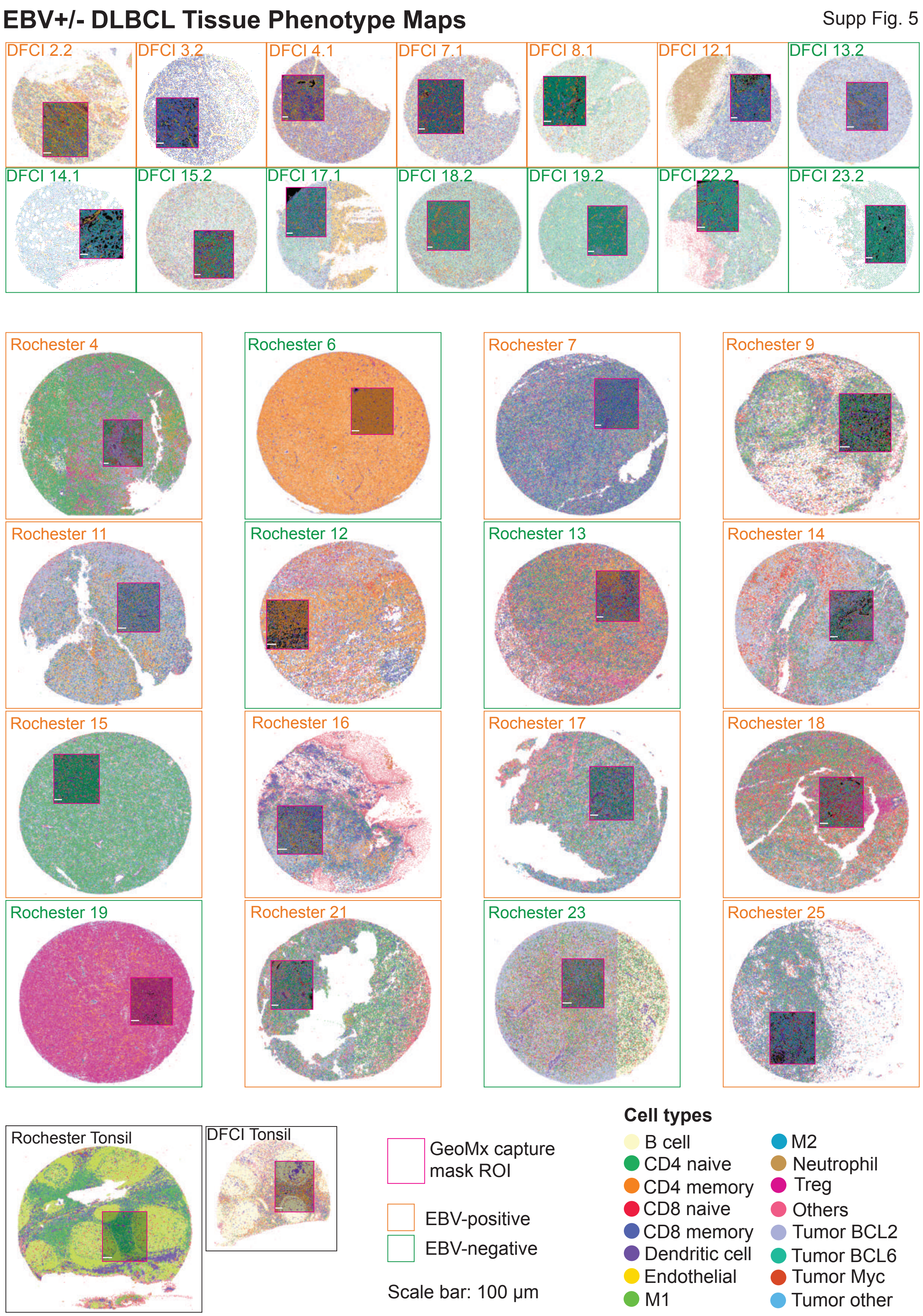


**Supplementary Figure 5: Validation of CODEX cell phenotyping.** Phenotype maps of each CODEX sample generated through iterative clustering and annotation based on single-cell CODEX features. Note that the DFCI cores (1.5mm diameter) are smaller than the Rochester cores (2.0mm diameter).

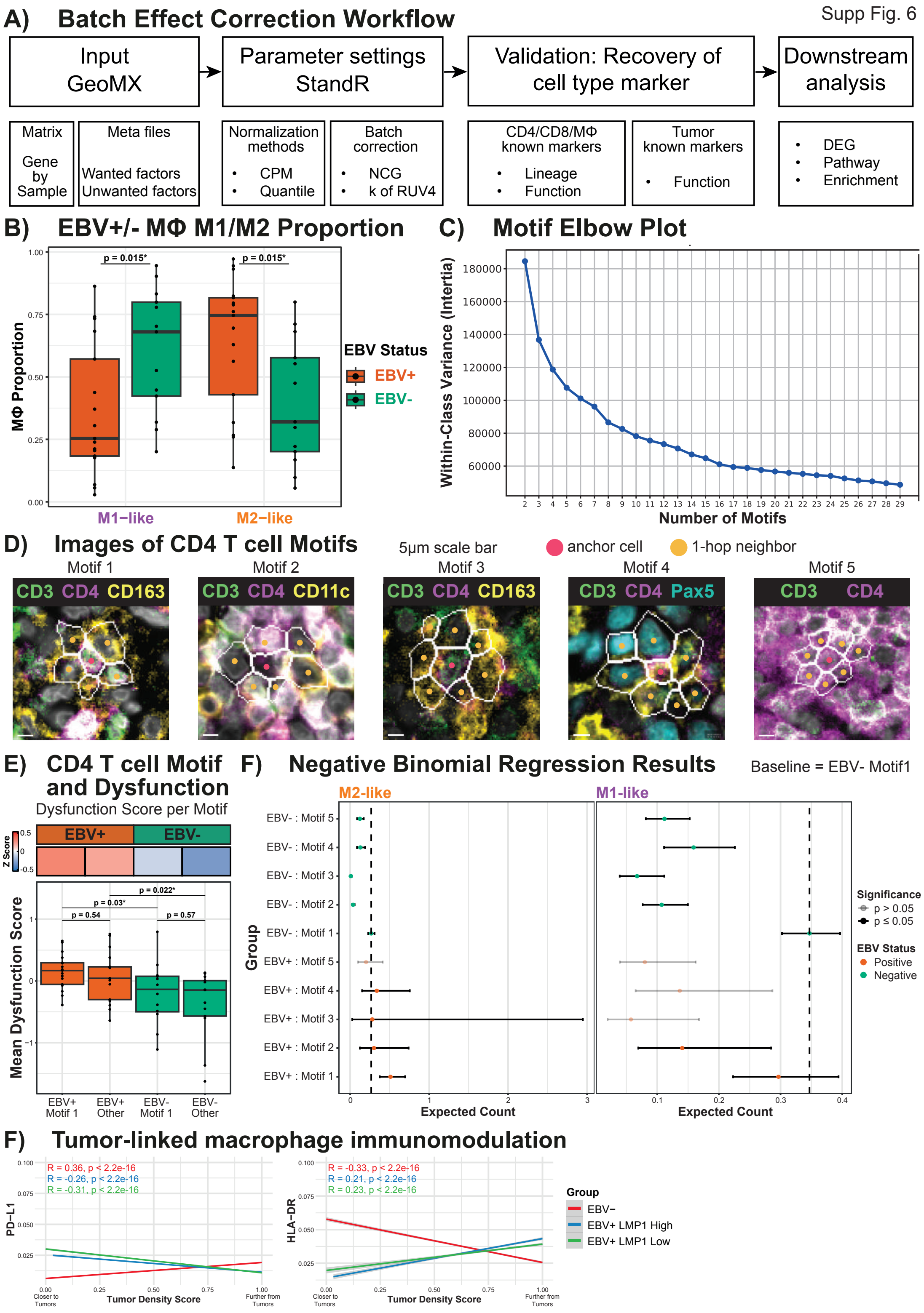


**Supplementary Figure 6: Evaluation of macrophage significance and optimal number of cellular motifs.** **(A)** Schematic of batch effect correction workflow. Detailed descriptions are in the **Materials and Methods** section. **(B)** Comparison of M1-like and M2-like macrophage proportions between EBV-positive and EBV-negative DLBCL tissues in this patient cohort, with each data point representing a sample. A two-sided Wilcoxon rank sum test was performed, with the null hypothesis that there is no difference in proportions between each respective M1-like or M2-like macrophages in EBV positive and that in EBV negative. **(C)** Elbow plot showing change in within-class variance with an increasing number of clustered motifs. **(D)** Representative immunofluorescence images of each motif showing CD4 T cells (CD4), tumor cells (Pax5), and macrophages (CD68 and CD163), with the corresponding cell phenotype maps. (E) Heatmap (top) and boxplot (bottom) of the protein-level CD4 T-cell dysfunction score between immune-enriched motif 1 and all other motifs combined. The "Other" category was obtained by combining Motifs 2 + 3 + 4 + 5. **(E)** Left: Negative binomial regression results with the response variable being the count of M2-like macrophages. Right: Negative binomial regression results with the response variable being the count of M1-like macrophages. For both regressions, the independent variables are EBV status, membership of motifs, and the interaction between them. The Y axis shows each independent variable and the X axis shows the fold change derived from exponentiating the coefficient of the independent variables (log fold change). For each variable, the point estimate is represented by a solid dot, with its corresponding 95% confidence interval shown as the bars surrounding the dot. **(F)** Correlation of PD-L1 and HLA-DR expression on macrophages with tumor cells stratified by LMP1 expression status. Details about tumor density score calculation are in **Materials and Methods**.

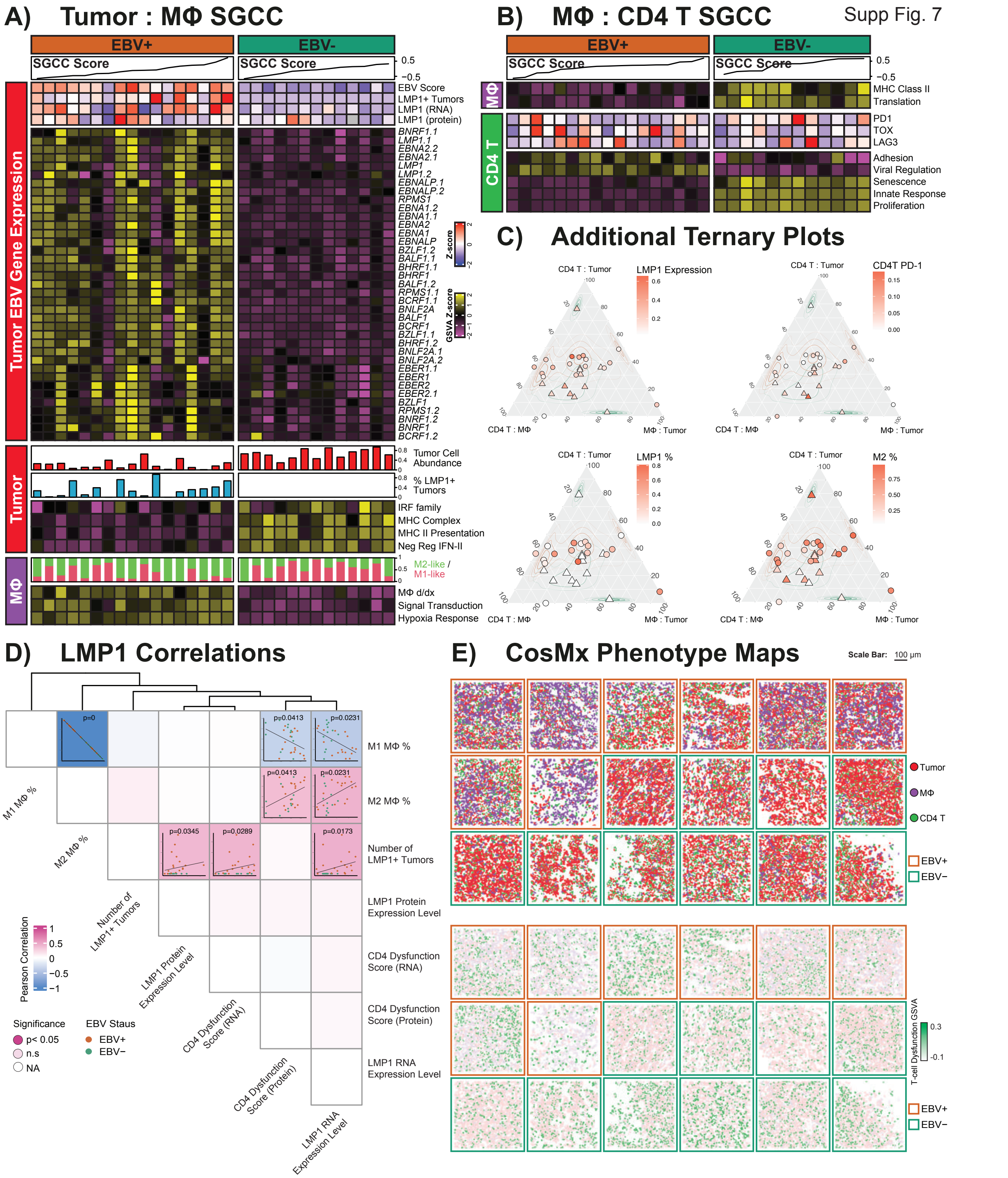


**Supplementary Figure 7: Additional supporting data for SGCC analysis of tumor-macrophage-CD4 T cell axis. (A)** Top: Expression of LMP1 and EBV transcripts in each ROI. Middle and Bottom: Additional data supporting contrasting M1/M2 polarization in EBV-negative/EBV-positive DLBCL. **(B)** Additional data supporting contrasting CD4 T-cell functional states between EBV-positive and EBV-negative DLBCL. **(C)** Additional ternary plot showing LMP1 expression on tumor cells, PD-1 on CD4 T cells, and proportion of M2-like macrophages. **(D)** Correlation of LMP1-expressing tumors with CD4 T-cell dysfunction signatures and macrophage M1/M2 polarization state across protein and RNA levels. **(E)** Top: Phenotype map of all CosMx regions showing the spatial organization of annotated tumor (red), macrophage (purple), and T-cell populations (green). Bottom: The same phenotype map colored to emphasize T-cell dysfunction signature on every annotated T cell.
